# Physics-Guided Generative Modeling Reveals that Steric Conflict Resolution Drives Ligand Specificity and Induced-Fit Adaptation

**DOI:** 10.64898/2026.11.28.769850

**Authors:** Diya Zhang, Chengwei Ai, Yuhao Liu, Xiaoyi Liu, Shiqiang Ma, Cheng Liang, Housheng Su, Shuangjia Zheng, Qiaozhen Meng, Fei Guo

## Abstract

Protein–ligand recognition is governed by the mutual resolution of steric conflicts as protein and ligand establish compatible interactions within a shared binding environment. Local chemical differences alter intermolecular interactions and influence both ligand discrimination and the conformational rearrangements required to achieve a physically compatible bound state. However, how steric conflict resolution links ligand discrimination to coupled conformational adaptation remains incompletely understood. Here, we develop CoupleDiff, a physics-guided generative approach for coupled protein–ligand conformational generation. Lennard–Jones interaction priors are incorporated into cross-molecular attention to distinguish repulsive, near-equilibrium, and attractive contact regimes, while geometric and force-based constraints guide the reverse diffusion process through the progressive resolution of intermolecular steric conflicts. Across six cross-docking datasets, CoupleDiff achieves the highest docking success rate, improving by 9.8% over the strongest competing method, together with the lowest RMSD among the evaluated methods, while reducing unfavorable intermolecular contacts. In alpha-1-acid glycoprotein 2, comparison of the structurally similar ligands disopyramide and isopropamide reveals how a single-atom chemical difference alters local interaction patterns, including a stabilizing hydrogen bond with Arg90, and determines binding-site compatibility. In carbonic anhydrase II, CoupleDiff recovers ligand-associated rearrangements of HIS200 and TRP209 from the apo structure, accompanied by conformational adjustment of the ligand. Molecular dynamics simulations further support the stability of the generated complex, while steric conflicts progressively decrease during reverse diffusion. Steric conflict resolution thus provides a physical link between ligand discrimination and coupled conformational adaptation, offering insight into the physical basis of protein–ligand recognition.

**Significance:** Protein–ligand recognition requires molecular partners to mutually resolve steric conflicts while establishing compatible interactions within a shared binding environment. How this process contributes to ligand discrimination and conformational adaptation remains incompletely understood. Using physics-guided generative modeling, we examine how local chemical differences alter the resolution of intermolecular steric conflicts between related ligands and how coordinated protein and ligand rearrangements accommodate the binding environment. Our results connect two important aspects of molecular recognition: distinguishing between chemically related molecules and adapting molecular conformations through the resolution of steric conflicts. This work provides a physical perspective on how steric conflict resolution underlies protein–ligand recognition and offers a computational approach for investigating and designing molecular interactions.

---

A single intermolecular contact can influence whether a ligand adopts a physically plausible binding mode, but molecular recognition depends on how a protein and a ligand jointly resolve steric conflicts while establishing compatible interactions within a shared binding environment (1, 2). As the two molecules interact, steric exclusion and short-range repulsion restrict their accessible conformations, while favorable noncovalent interactions contribute to the stabilization of compatible binding arrangements. Binding therefore involves more than geometric accommodation of a ligand within a pocket; it requires the coordinated satisfaction of local chemical and structural constraints by both molecular partners. Local steric conflicts and their resolution can influence ligand selectivity, binding-site accommodation, and the conformational changes associated with complex formation. Understanding the physical basis of intermolecular compatibility is consequently important for explaining molecular recognition and advancing structure-based drug discovery (SBDD) (3–5).

Protein–ligand recognition involves multiple physical factors, including solvation, entropy, electrostatics, and receptor pre-organization. Among these factors, local intermolecular steric and interaction constraints provide a direct structural perspective on how binding configurations are excluded or accommodated (6, 7). Local chemical and geometric incompatibilities can have distinct consequences depending on whether the interacting molecules can accommodate them. Persistent incompatibilities can disfavor particular binding configurations and contribute to ligand discrimination, even between structurally similar molecules (8–12). Alternatively, local incompatibilities can be accommodated through side-chain reorientation, binding-pocket rearrangement, or ligand torsional adjustment, allowing protein and ligand to adopt mutually compatible conformations (13–15). Ligand discrimination and conformational adaptation therefore represent two related structural outcomes of intermolecular physical compatibility: unfavorable interactions can restrict molecular recognition, whereas conformational rearrangements can relieve local conflicts and enable binding (16, 17).

Coupled conformational adaptation also raises a long-standing question in molecular recognition: whether a ligand selects a receptor conformation already present in solution or promotes the formation of a new conformation upon binding (18–21). Static complex structures capture the final arrangement of interacting molecules but provide limited information about the structural adjustments leading to the bound state. Structural comparisons and molecular simulations offer complementary information about conformational preferences and accessible states, yet connecting local intermolecular interactions to coordinated receptor and ligand rearrangements remains challenging. In particular, understanding how the resolution of local steric conflicts relates to the conformational changes accompanying ligand binding requires consideration of both molecular partners and their interaction environment. A description of protein–ligand recognition that connects local interaction compatibility with coupled conformational adaptation could therefore provide insight beyond the structural characterization of final bound states.

Computational approaches differ in how intermolecular physical constraints are represented during complex structure prediction. Classical docking programs, including AutoDock, Glide, and GOLD, incorporate steric and van der Waals (vdW) terms into their scoring functions, but generally evaluate ligand poses against fixed or only partially flexible receptors, limiting their ability to capture coupled protein–ligand rearrangements (22–26). Learning-based methods capture increasingly complex structural and interaction patterns: FeatureDock, FlexPose, and Interformer learn interaction geometry from structural data (27–29), while interaction-aware models improve affinity prediction and generalization (30). Generative approaches further extend conformational sampling to receptor flexibility and physical information. DynamicBind models receptor motion associated with ligand binding (14), DiffBindFR jointly samples ligand and side-chain conformations (31), and physics- or surface-informed diffusion approaches incorporate force or interaction information into structure generation (32–34). Despite advances in flexible conformational sampling and physical modeling, steric clashes and distorted local geometries remain important challenges in generated complexes (35, 36). More fundamentally, treating steric clashes primarily as structural violations to be penalized or filtered does not explicitly represent the physical process through which protein and ligand resolve local conflicts during conformational generation. Recovering a native-like bound structure alone also provides limited insight into how local interaction differences contribute to ligand discrimination or how conformational rearrangements accommodate intermolecular incompatibilities.

Here, we develop CoupleDiff, a physics-guided generative approach for coupled protein–ligand conformational generation that explicitly incorporates intermolecular steric conflict resolution into the generation process. Lennard–Jones (LJ) interaction priors modulate cross-molecular attention, enabling the model to distinguish repulsive, near-equilibrium, and attractive contact regimes when representing intermolecular interactions. Geometric and force-based constraints further guide the reverse diffusion process through the progressive resolution of steric conflicts, directing the joint generation of protein and ligand conformations toward physically compatible structures. Across six cross-docking datasets, CoupleDiff achieves the highest docking success rate and the lowest RMSD among the evaluated methods, while reducing unfavorable intermolecular contacts. Ablation analyses further assess the contributions of interaction-informed attention and constraint-guided conformational generation (SI Appendix C). Beyond pose recovery, we examine two complementary molecular systems to investigate how local interaction differences and steric conflict resolution relate to distinct aspects of molecular recognition. In alpha-1-acid glycoprotein 2 (AAGP2), comparison of the structurally similar ligands disopyramide and isopropamide illustrates how a single-atom chemical difference alters local interaction patterns, including a stabilizing hydrogen bond with Arg90, and influences binding-site compatibility. In carbonic anhydrase II (CAII), CoupleDiff recovers ligand-associated rearrangements of HIS200 and TRP209 from the apo receptor structure, accompanied by conformational adjustment of the ligand. Molecular dynamics simulations further support the stability of the generated complex. The results connect local chemical differences with ligand discrimination and the progressive resolution of intermolecular steric conflicts with coupled conformational adaptation, providing a physical perspective on protein–ligand recognition.

## Results

### A physics-guided framework that resolves conflicts during generation

Protein–ligand binding occurs within a spatially and chemically constrained environment, where steric exclusion and intermolecular interactions determine whether a candidate binding state is physically accessible. These constraints must therefore be resolved during the formation of a binding complex rather than evaluated only after a pose has been generated. Guided by this principle, CoupleDiff was developed to incorporate physical constraints directly into protein–ligand conformational generation (Fig. 1).

**Fig. 1.**
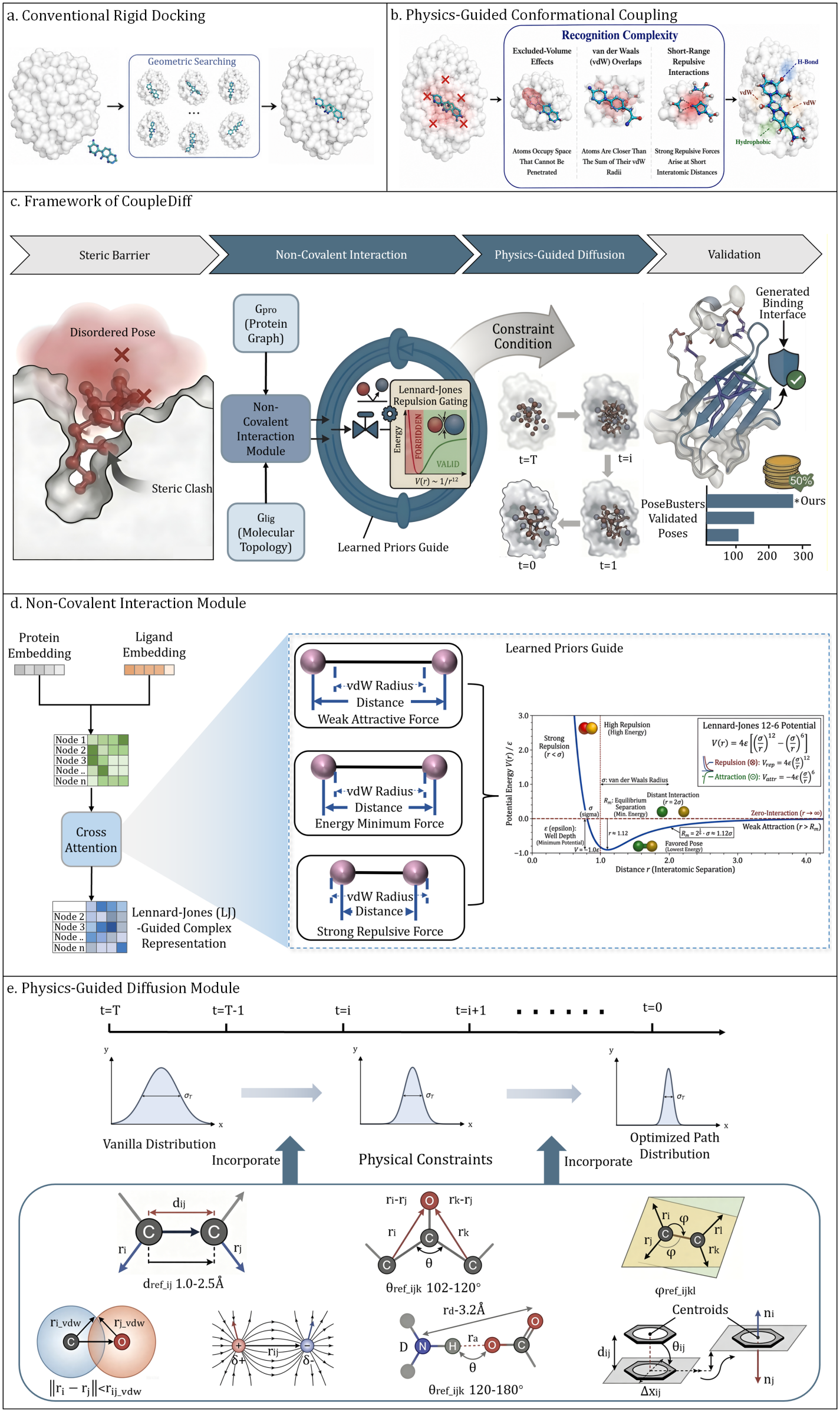
Overview of CoupleDiff, a physics-guided diffusion framework for protein–ligand conformational generation. **a** Protein and ligand conformations are typically treated as constrained during the conventional pose generation, with the search focused on identifying geometrically favorable ligand poses. **b** Starting from an unfavorable initial pose, CoupleDiff progressively resolves physical incompatibilities through interaction-aware representation learning and physics-guided diffusion, enabling coordinated adaptation of protein and ligand conformations toward a physically compatible binding state. **c** CoupleDiff incorporates physical constraints at both the representation and conformational generation levels. Protein and ligand representations are first coupled through a non-covalent interaction module, in which LJ-guided gating encodes intermolecular physical compatibility. The resulting conditional representations then guide a physics-constrained diffusion process for iterative protein–ligand pose refinement. Generated complexes are subsequently evaluated for structural and physical plausibility. **d** Protein and ligand embeddings are fused through cross-attention, where attention scores are dynamically modulated by LJ interaction priors, to distinguish sterically repulsive, near-equilibrium, and weakly attractive regimes. **e** Protein–ligand pose generation is formulated as a constrained diffusion process. During reverse diffusion, physical force and geometric constraints reshape the denoising trajectory and guide the coupled conformational evolution toward physically compatible binding states.

CoupleDiff formulates protein–ligand binding as a coupled conformational generation process, in which protein and ligand conformations are jointly refined under physical interaction and geometric constraints (Fig. 1b), unlike conventional docking approaches that search for ligand poses within a predefined receptor conformation (Fig. 1a). The framework consists of two complementary components: a non-covalent interaction module and a physics-guided diffusion module (Fig. 1c).

The generation process begins with protein graph representations (*G*_pro_) and ligand topology embeddings (*G*_lig_), which are integrated through a non-covalent interaction module (Fig. 1d). This module enables bidirectional information exchange between protein and ligand atoms, allowing the model to capture cross-molecular dependencies that govern binding. Rather than relying solely on learned attention weights, CoupleDiff introduces interaction priors derived from intermolecular distances to distinguish favorable contacts from unfavorable contacts. Interatomic distances are projected onto the LJ interaction, partitioning contacts into repulsive, equilibrium, and weakly attractive regimes. These interaction states dynamically modulate cross-attention scores, suppressing unfavorable contacts while enhancing physically compatible interactions. As a result, the learned representations encode not only geometric proximity but also physically informed intermolecular relationships.

The resulting conditional embeddings are subsequently used to guide a constrained diffusion process (Fig. 1e). During reverse diffusion, protein and ligand coordinates are iteratively refined under complementary intramolecular and intermolecular physical constraints, including bond lengths, bond angles, and torsions, vdW exclusion, electrostatics, hydrogen bonding, and *π*–*π* stacking. These constraints reshape the denoising trajectory and guide sampling toward conformations that satisfy coupled geometric and interaction requirements.

### Explicit physical constraints reduce steric clashes and recover native interactions

A physically meaningful binding state must reproduce not only the overall geometry of a protein– ligand complex, but also the specific non-covalent interactions that stabilize molecular recognition. We first evaluated the ability of the predicted complexes to recover interaction patterns observed in experimental structures. Across five interaction types, CoupleDiff recovered 59% of hydrogen bonds, 40% of hydrophobic interactions, 77% of salt bridges, 31% of *π*-stacking interactions, and 35% of *π*-cation interactions (Fig. 2a). CoupleDiff achieved the highest recovery rate across all five interactions, with pronounced improvements in salt-bridge and *π*-mediated interactions. These results indicate that the structural predictions are accompanied by more accurate reconstruction of interaction networks, extending beyond the recovery of global geometric similarity.

**Fig. 2.**
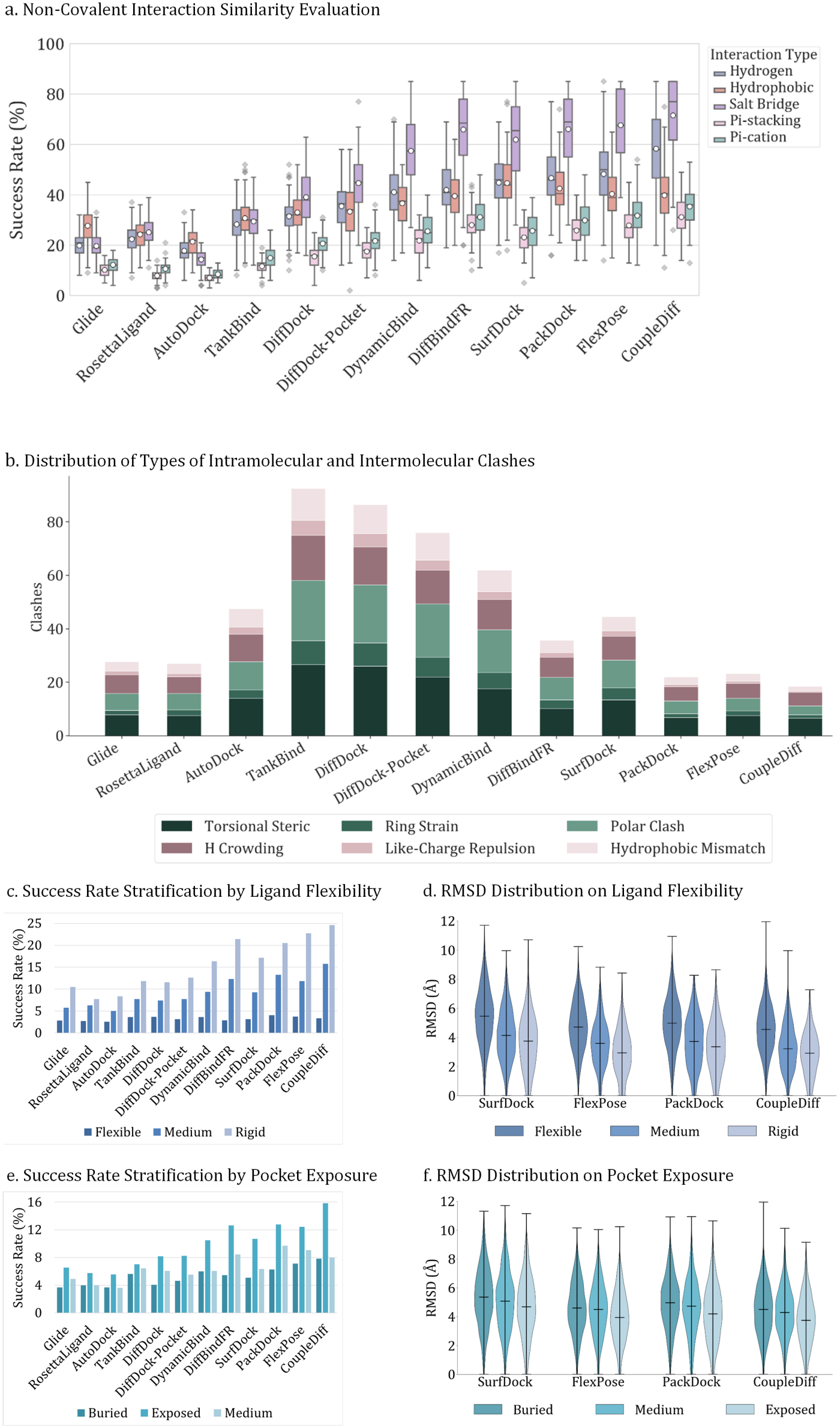
Explicit physical constraints improve the physical feasibility of predicted binding states. **a** Recovery of five major non-covalent interaction types in predicted protein–ligand complexes. **b** Distribution and composition of physical conflicts in generated complexes. Overall conflict counts are compared across methods, with conflicts further categorized into torsional steric conflicts, ring strain, polar clashes, hydrogen crowding, like-charge repulsion, and hydrophobic mismatches. **c, d** Success rate and RMSD stratified by ligand rotatable bond counts, assessing robustness to increasing ligand conformational freedom. **e, f** Success rate and RMSD stratified by pocket exposure, assessing robustness across binding sites with different degrees of spatial confinement.

However, accurate interaction recovery does not necessarily guarantee that a predicted complex is physically realizable. We therefore assessed the geometric integrity of the generated structures by quantifying both intramolecular conflicts within the ligand and intermolecular conflicts between the ligand and protein on the PoseBusters(PB) benchmark (Fig. 2b). Classical docking approaches generally produced fewer steric conflicts, consistent with their treatment of steric exclusion during pose optimization. In contrast, several deep generative approaches such as TankBind (37), DiffDock (38), DiffDock-Pocket (39), and DynamicBind (14) exhibited broader distributions and more severe outliers in both intramolecular and intermolecular conflicts. CoupleDiff showed the lowest median conflict counts for both intramolecular and intermolecular clashes, together with a substantially narrower distribution across the benchmark.

To determine whether the reduction in physical conflicts was restricted to a particular type of structural violation, we analyzed six major categories of conflicts: torsional steric conflicts, ring strain, polar clashes, hydrogen crowding, like-charge repulsion, and hydrophobic mismatches (Fig. 2b). Several competing generative methods showed higher conflict counts in multiple categories, with torsional steric conflicts and polar clashes being the most common. In contrast, CoupleDiff consistently reduced violations across all six categories, indicating a general improvement in physical consistency rather than correction of a single failure mode.

We next examined whether the benefit of explicit physical constraints persists as the conformational search space becomes increasingly complex. Ligands were stratified according to the number of rotatable bonds into rigid (*<*5), intermediate (5–10), and flexible (*>*10) groups. As expected, most methods showed reduced docking success with increasing ligand flexibility (Fig. 2c,d). CoupleDiff exhibited a substantially smaller performance decrease and retained the highest success rate in the most flexible group. The corresponding RMSD distributions showed a similar trend, indicating that the advantage of physical constraint-aware generation is maintained as ligand conformational freedom increases.

We further evaluated whether the benefit of physical constraint modeling depends on the spatial accessibility of the binding site. Binding pockets were classified as buried, intermediate, or exposed according to their solvent exposure (Fig. 2e,f). Most methods showed reduced success in buried pockets, where ligand accommodation occurs under stronger geometric confinement. CoupleDiff maintained the highest success rate across all three exposure regimes and showed a particularly pronounced advantage for buried binding sites. Taken together, these results show that physical feasibility can be built into conformational generation itself, and that this advantage holds as ligand flexibility and pocket burial increase.

### CoupleDiff recovers native poses under receptor conformational heterogeneity

In realistic SBDD, the receptor conformation that matches a productive binding state is rarely available as a single, pre-organized structure. Docking may begin from apo receptors, non-cognate holo structures, or imperfect homology models, where the binding environment differs from the native complex. This heterogeneity poses a basic challenge: the model must find a productive ligand pose from a receptor geometry that does not present the native binding environment. We evaluated CoupleDiff in cross-docking settings on four datasets that span these conditions, CDK2, EGFR, DUDE27-HoloEns, and ANN, which range from local induced-fit changes to large backbone and side-chain perturbations; two further cross-docking datasets, CASF2016-Drug and a PDBbind cross-docked core set, are reported in SI Appendix 3.B. As a reference, we also evaluated redocking from native-like receptors, where the binding environment is already pre-organized (SI Appendix 3.A). Across the cross-docking datasets, CoupleDiff achieved the highest success rate and the lowest median RMSD, raising the mean success rate from 48.1% for the strongest prior method to 57.9% (SI Appendix, Table S6).

We first examined apo-to-holo settings, where binding requires the ligand to fit a receptor that differs from the experimental complex. On CDK2, CoupleDiff produced a higher fraction of near-native poses (Fig. 3a) and the lowest median RMSD among the evaluated methods (Fig. 3b). EGFR dataset showed a similar pattern, where differences in the activation loop and *α*C-helix add further conformational challenges (Fig. 3c, d). Recovering nearnative conformations from these non-native receptors shows that physics-guided generation stays effective when the starting receptor does not provide the geometry of the productive binding state, and the predicted complexes move closer to the experimental binding state even when the input receptor is not pre-organized.

**Fig. 3.**
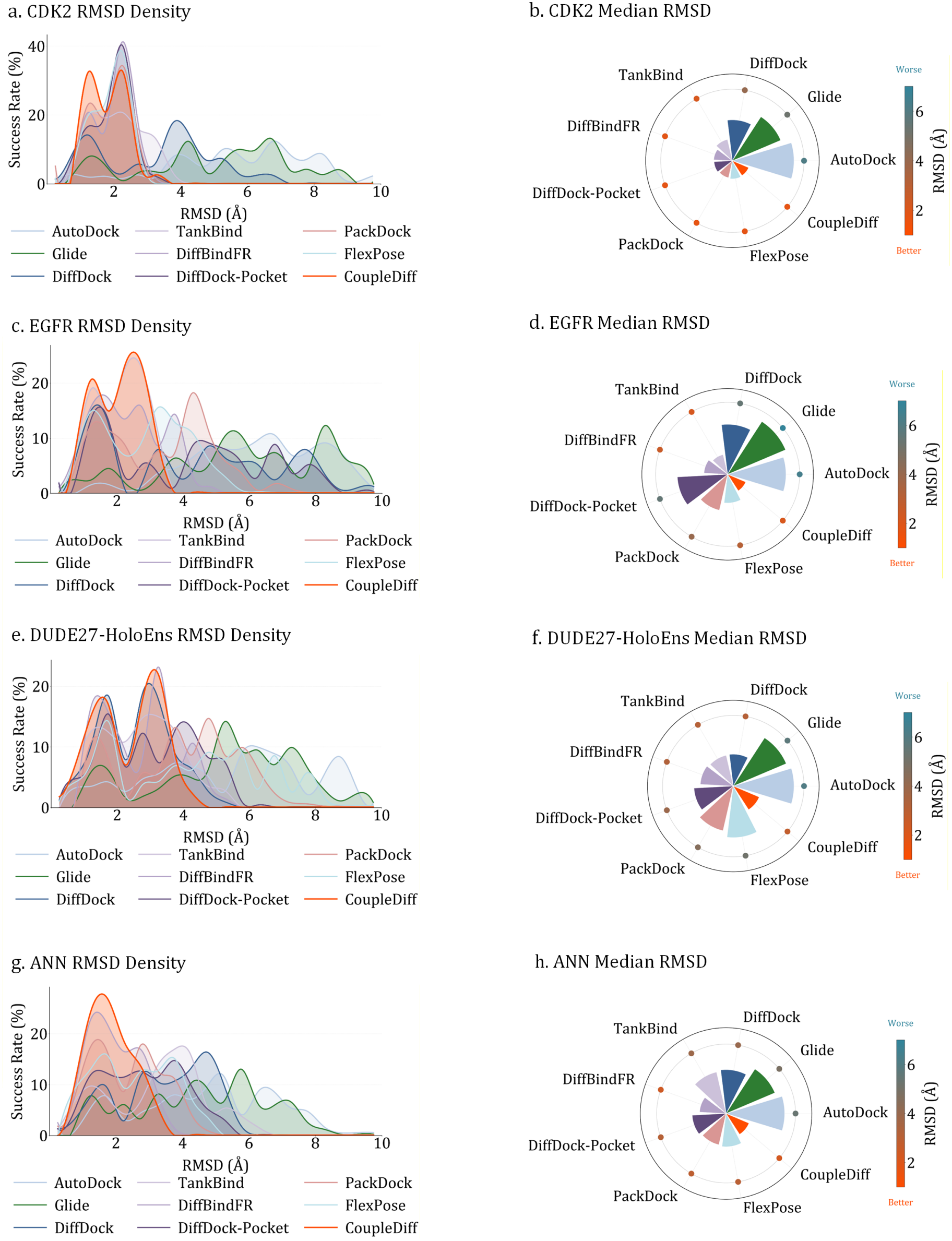
Cross-docking evaluation across four benchmarks spanning distinct forms of receptor conformational heterogeneity. **a, b** CDK2, representing apo-to-holo receptor adaptation. **c, d** EGFR, representing conformational changes between apo and holo states. **e, f** DUDE27-HoloEns, representing holo-to-holo cross-docking. **g, h** ANN, representing docking against structurally perturbed non-native receptor models. For each benchmark, pose distributions across RMSD values and corresponding median RMSD comparisons are shown.

We next considered holo-to-holo cross-docking, where the receptor is already organized around a different, non-cognate ligand. The starting receptor provides a pocket, but one shaped by another ligand and possibly incompatible with the ligand being docked. On DUDE27-HoloEns, CoupleDiff shifted the RMSD distributions toward the near-native regime (Fig. 3e) and reached the lowest median RMSD among the compared methods (Fig. 3f). Recovering near-native poses from pockets pre-organized by other ligands shows that the model generalizes across receptor conformations produced by different ligand environments, beyond the geometry of any single ligand-bound template.

Finally, the ANN dataset introduces large uncertainty through homology models and unrelated non-native receptor structures, with backbone displacements, inaccurate side chains, and altered loop geometries that can differ markedly from the native complex. Despite these perturbations, CoupleDiff kept a high precision of near-native predictions (Fig. 3g) and the lowest median RMSD among the evaluated methods (Fig. 3h). Recovering near-native conformations from such heterogeneous receptors shows that the model can find interaction-compatible binding states without a starting receptor that closely resembles the native complex.

Taken together, across apo-to-holo, holo-to-holo, and perturbed homology-model settings, CoupleDiff recovered near-native conformations despite large differences between the starting receptor and the experimental complex, which proves that building these constraints into generation improves the recovery of binding states across heterogeneous receptors.

### CoupleDiff distinguishes near-identical ligands at AAGP2

Molecular recognition ultimately requires discrimination between chemically related molecules that can occupy similar spatial regions but differ in their ability to form stabilizing interactions. This provides a more stringent test of physical constraint-guided recognition than recovery of a single native binding pose. To examine whether CoupleDiff can resolve such subtle molecular differences, a chemically controlled case study was designed using human AAGP2 (40) and a pair of structurally similar drugs with divergent binding behavior. The two compounds share a highly similar scaffold and were evaluated at the same protein binding site, thereby minimizing differences in overall molecular geometry while exposing the contribution of local chemical complementarity.

Disopyramide, an antiarrhythmic drug with experimental binding to AAGP2, was used as the positive control (41). To identify a matched negative control, approved small-molecule drugs from DrugBank were compared using extended-connectivity fingerprints and Tanimoto similarity coefficients. t-distributed stochastic neighbor embedding(t-SNE) visualization of the chemical space identified isopropamide as a close structural analogue of disopyramide, with a Tanimoto similarity of 0.92 (Fig. 4i). Despite this high structural similarity, isopropamide is not a known AAGP2 ligand and therefore provides a controlled comparison for molecular discrimination.

**Fig. 4.**
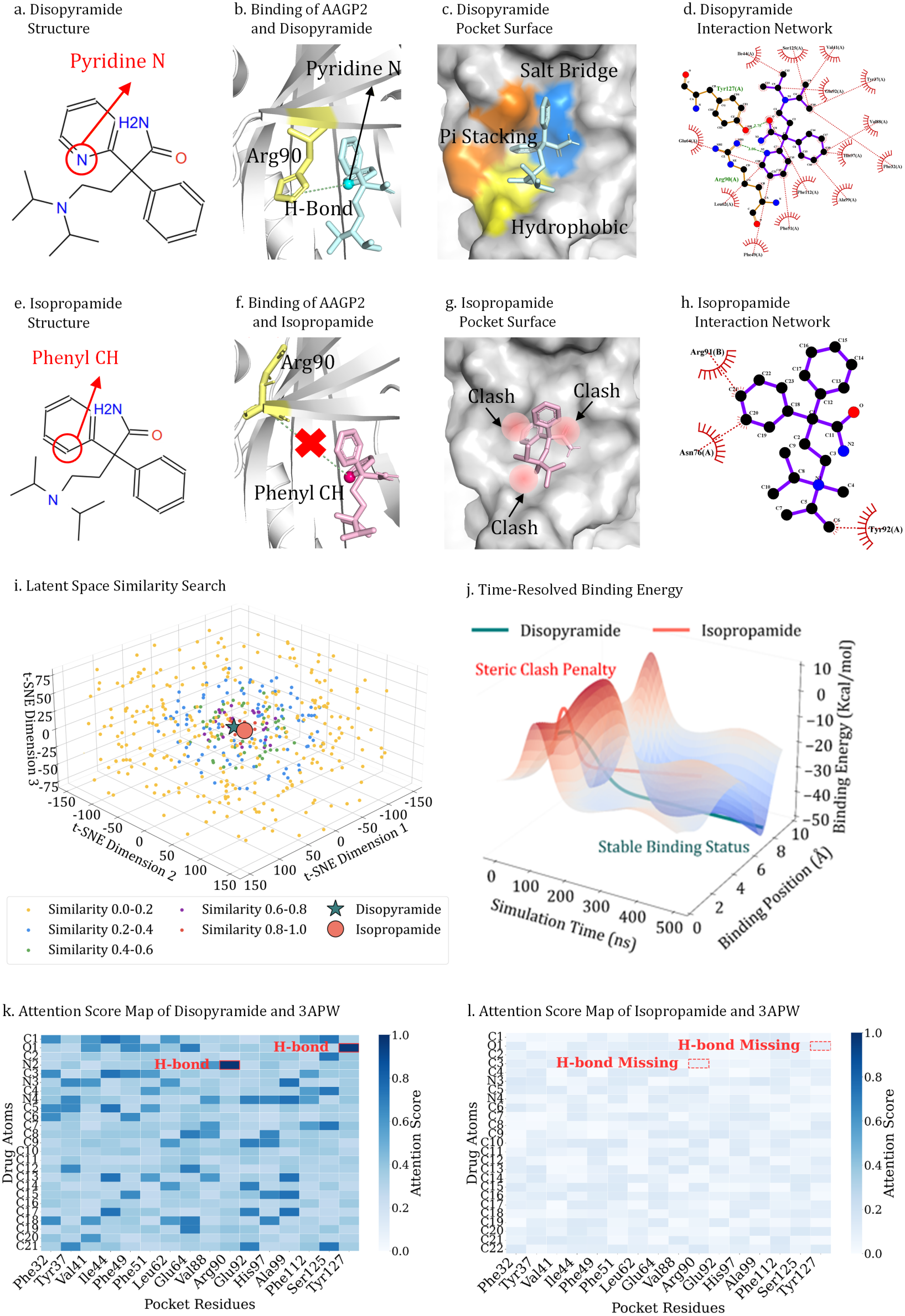
Physical constraints enable molecular discrimination between near-identical analogs at the AAGP2 target. **a** Disopyramide structure highlighting the pyridine nitrogen that serves as a hydrogen-bond acceptor. **b** Predicted AAGP2–disopyramide complex showing the hydrogen bond between the pyridine nitrogen and Arg90. **c** Disopyramide binding pocket showing cavity formation and favorable surrounding contacts. **d** Two-dimensional interaction map showing the stabilizing interaction network formed by disopyramide. **e** Isopropamide structure showing substitution of the pyridine nitrogen with a phenyl group. **f** Predicted AAGP2–isopropamide complex showing loss of the hydrogen-bonding interaction and unfavorable proximity between the phenyl group and Arg90. **g** Isopropamide binding environment showing regions of ligand–protein physical incompatibility. **h** Two-dimensional interaction map showing reduced stabilizing interactions and increased unfavorable contacts for isopropamide. **i** Chemical similarity landscape identifying isopropamide as the closest structural analogue of disopyramide. **j** Time-resolved interaction-energy profiles of disopyramide and isopropamide in complex with AAGP2. **k, l** Attention score maps for AAGP2 complexes with disopyramide and isopropamide, respectively.

The two compounds exhibited markedly different physical behavior in the AAGP2 binding environment. Disopyramide rapidly reached a low and stable interaction energy, consistent with formation of a favorable bound state, whereas isopropamide maintained higher and more variable interaction energies without reaching a stable regime (Fig. 4j). The change lies in the aromatic ring beside the shared core: disopyramide carries a pyridine whose nitrogen can act as a directional hydrogen-bond acceptor, while isopropamide carries a phenyl ring that cannot (Fig. 4a, e).

This small chemical difference produces a distinct local interaction environment at the AAGP2 binding site. In the predicted disopyramide complex, the pyridine nitrogen is oriented toward Arg90 and forms a hydrogen bond that contributes to anchoring the ligand within the pocket (Fig. 4b). Isopropamide occupies a similar spatial region but lacks the corresponding hydrogen-bond acceptor. Its phenyl group instead approaches Arg90 without providing a compatible polar interaction, producing an unfavorable local arrangement (Fig. 4f).

The difference extends beyond the single Arg90 interaction. Disopyramide is accommodated within a defined cavity containing additional hydrophobic, *π*–*π* stacking, and electrostatic interactions (Fig. 4c). In contrast, the isopropamide complex exhibits multiple regions of ligand–protein overlap and lacks an equivalently coherent interaction network (Fig. 4g). Consistent with these structural observations, the corresponding two-dimensional interaction maps show a dense network of favorable contacts for disopyramide, whereas isopropamide displays fewer stabilizing interactions and more unfavorable contacts (Fig. 4d, h).

The molecular distinction is also reflected in CoupleDiff’s interaction representations. The attention map for disopyramide contains localized high-attention regions at chemically meaningful interaction sites, including the pyridine nitrogen–Arg90 pair and additional contacts involving Tyr127 and hydrophobic residues such as Phe49 and Phe112 (Fig. 4k). In contrast, the isopropamide complex exhibits a more diffuse attention pattern and lacks comparable focal interactions(Fig. 4l). The correspondence between attention hotspots and interaction sites indicates that the model representation captures local chemical features relevant to recognition, and not only global shape similarity.

At AAGP2, the loss of this hydrogen-bond acceptor propagates from local interaction to steric accommodation and overall binding organization. With the benchmark analyses above, these results show that physical constraints support molecular discrimination: small chemical differences translate into physically compatible or incompatible binding states that the model can tell apart.

### CoupleDiff Predicts the Induced-Fit Pocket of Carbonic Anhydrase

**II**. Molecular recognition can require the receptor to reorganize its binding site in response to the incoming ligand. A physics-guided model should reproduce such reorganization where it occurs. We tested this on CAII and its inhibitor acetazolamide, providing the apo receptor as input and asking whether CoupleDiff generates the ligand-bound pocket.

The conformational response in CAII is concentrated at the active site. The protein backbone differs by only *∼* 0.75 Å between the apo and holo states, yet across the pocket residues (HIS200, TRP209, LEU158, THR190, and HIS119) the heavy-atom conformation shifts by 1.87 Å, driven mainly by reorientation of the HIS200 and TRP209 side chains that enlarges the space available to the ligand (Fig. 5a). Starting from the apo structure, CoupleDiff reproduced the holo pocket to within 0.79 Å heavy-atom RMSD, below the 1.87 Å separating apo from holo, while moving 2.23 Å away from its apo input.

**Fig. 5.**
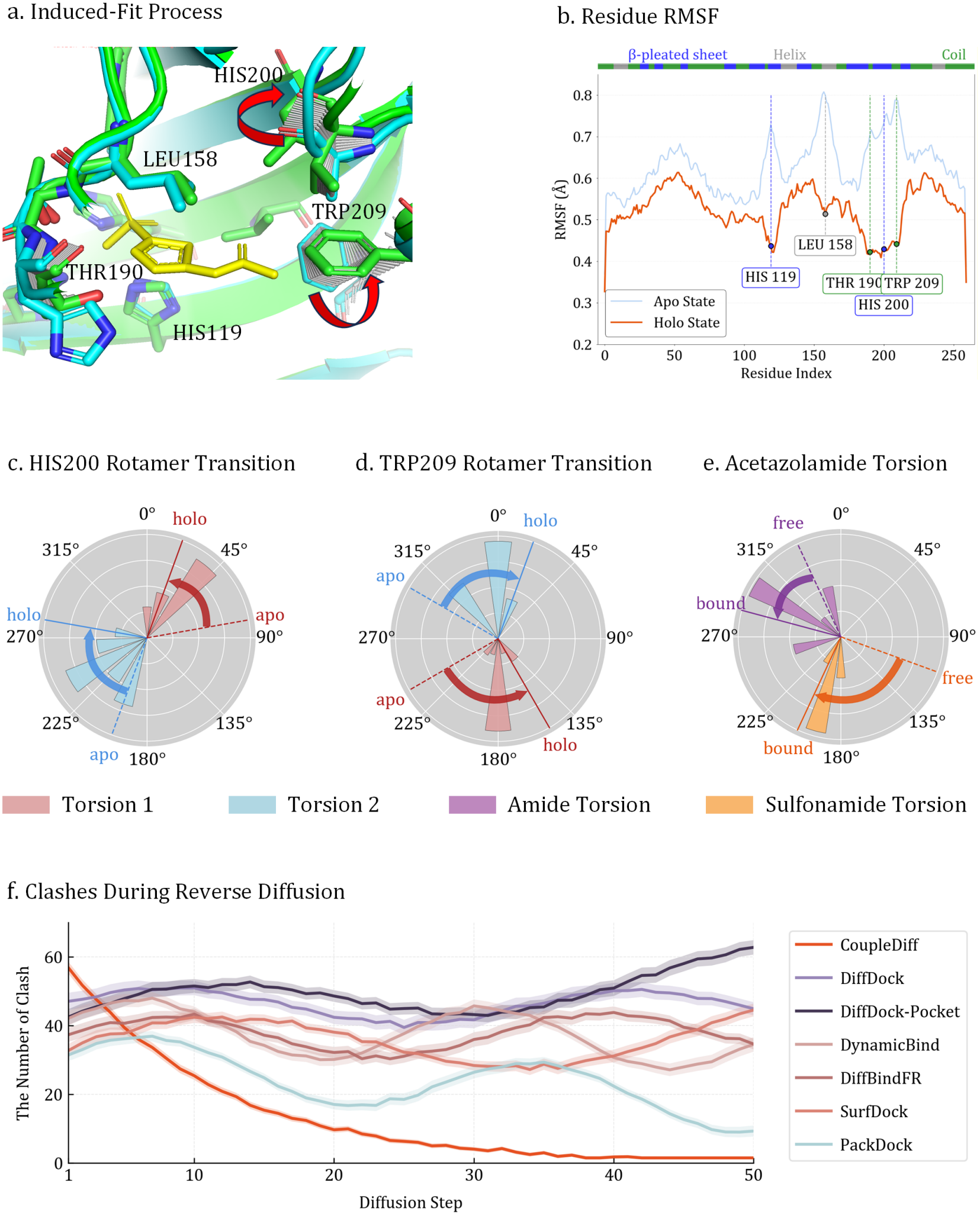
CoupleDiff predicts the induced-fit pocket of CAII. **a** Structural comparison of apo (cyan) and holo (green) CAII states. Acetazolamide is shown as yellow sticks. Red arrows indicate the side-chain reorientations of HIS200 and TRP209 that expand the accessible space of the sterically occluded binding pocket. **b** Residue-wise RMSF profiles of the CAII active site from MD simulations of the apo and holo states. **c** HIS200 rotamer transition: The shift in side-chain dihedral angles from the apo state to the holo-specific orientation. **d** TRP209 rotamer transition: The realignment of the indole ring to stabilize hydrophobic and pi-stacking interactions. **e** Acetazolamide torsion: The distribution of the amide and sulfonamide torsional angles. **f** Clashes during reverse diffusion: Evolution of steric clashes throughout the reverse diffusion process.

Two questions follow: whether this rearrangement is induced by binding rather than selected from a pre-existing ensemble, and whether the predicted complex is physically stable. Both were addressed by molecular dynamics(MD). Over 500 ns of apo CAII, HIS200 and TRP209 did not sample their holo rotamers (Fig. 5c,d): the bound side-chain conformation is not present in the apo ensemble and is therefore induced upon binding rather than selected from it.

By contrast, MD of the CoupleDiff-generated complex remained within the holo rotameric basins, with HIS200 positioning its imidazole toward the ligand and TRP209 orienting its indole to support hydrophobic and *π*–mediated contacts with acetazolamide. Residue-wise root mean square fluctuation(RMSF) showed that binding selectively reduces mobility around the pocket, more strongly in loop and coil regions than in constrained secondary-structure elements (Fig. 5b). Together these simulations confirm that the predicted induced-fit conformation is stable.

Acetazolamide undergoes a corresponding conformational adjustment to accommodate the adapted binding site. The ligand contains two key torsional degrees of freedom, involving the amide and sulfonamide groups, that determine the orientation of its hydrogen-bond donors and acceptors. MD simulations revealed distinct changes in the distributions of these torsional angles between the unbound and bound states (Fig. 5e). In the unbound state, the amide torsion predominantly sampled the 270°– 315° region, whereas the sulfonamide torsion occupied a broader range between 90° and 180°, reflecting substantial conformational freedom. Upon binding, the amide torsion shifted toward a more confined 180°–270° region, accompanied by a redistribution of the sulfonamide torsion toward 180 *−* 225°. These shifts indicate that acetazolamide preferentially adopts a more specific geometry in the bound state, positioning its chemically active groups to satisfy the spatial and interaction constraints of the remodeled CAII pocket.

To examine the role of explicit physical constraints in reaching these states, the evolution of steric conflicts was therefore monitored throughout the reverse diffusion process (Fig. 5f). At the early denoising stages, generated structures were highly perturbed and contained substantial numbers of steric conflicts. As denoising proceeded, CoupleDiff showed a continuous reduction in the number of conflicts, decreasing from approximately 60 initial conflicts to nearly zero at the end of generation. In contrast, competing diffusion-based approaches, including DiffDock (38), DiffDock-Pocket (39), DynamicBind (14),

DiffBindFR (31), SurfDock (33), and PackDock (34), retained substantially more conflicts during generation and more frequently converged to states containing steric violations. The physical objectives in CoupleDiff act during reverse diffusion itself, shaping the intermediate conformations from which the final binding state emerges.

Together, these results show that CoupleDiff reproduces an induced-fit binding state and resolves the physical conflicts that gate it. These constraints both enforce physical feasibility and recover the local conformational adaptation that accompanies recognition.

## Discussion

Protein–ligand recognition has traditionally been read as geometric complementarity. Our results support a different account, in which recognition is a physically constrained process: as a complex forms, steric and energetic conflicts arise and are progressively resolved. CoupleDiff makes this process explicit by resolving such conflicts step by step during generation, and this single view connects two phenomena that are usually treated separately. When the conflicts introduced by a ligand cannot be resolved, the binding state is disfavored, which produces specificity; at AAGP2, a subtle chemical change that the pocket cannot accommodate shifts predicted compatibility between two closely related drugs. When the conflicts can be resolved, receptor and ligand adjust together, which produces induced fit. At CAII, the model reorganizes active-site side chains from the apo toward the holo. Molecular specificity and conformational adaptation are therefore two outcomes of one mechanism, the resolution of physical constraints, and both emerge from modeling how a complex forms as a physical process. This behavior follows from where the physical information is applied. In CoupleDiff, the interaction priors and the physics and geometry objectives act on the reverse-diffusion trajectory itself, so each intermediate conformation is pushed toward a physically compatible state as the complex is built. The conflict trajectories show the effect directly: steric violations fall monotonically through denoising and reach near zero, while methods that assess physical plausibility only after a pose is produced retain more conflicts. Applying the constraints during generation also lets the protein pocket and the ligand move in response to each other as the complex is built, which is what makes the coupled side-chain and ligand rearrangement at CAII accessible to the model. Physical feasibility and conformational adaptation are thus produced together, and both are difficult to recover from geometric fitting alone.

As with physics-guided generative modeling more broadly, several challenges remain. Existing benchmarks capture only part of the diversity of protein–ligand systems encountered in practice, and performance on targets that differ substantially from the training distribution remains an open question for the field. Incorporating increasingly detailed physical descriptions also raises the computational demands of generation. Addressing these points will require continued development of both datasets and modeling strategies.

In summary, CoupleDiff models protein–ligand complexes with improved physical feasibility and conformational accuracy by resolving steric and energetic conflicts during generation, and in doing so it captures adaptive rearrangements and binding specificity that approaches focused on geometric complementarity tend to miss. Beyond improving pose prediction, it offers a mechanistic account of how recognition can discriminate between chemically similar ligands or adapt to accommodate them, and it positions physical constraint resolution as a general perspective for understanding and modeling protein–ligand recognition.

## Materials and Methods

### Details of CoupleDiff

CoupleDiff is a transformer-based diffusion model designed for accurate modeling of protein–ligand complexes, which is composed of three principal modules working in synergy. First, a representation encoding module utilizes a cooperative scalar-vector mechanism to generate tightly coupled geometric and chemical feature embeddings for the protein backbone, side chains and the ligand. The backbone embedding is further discretized into a semantic token space. Second, a non-covalent interaction module fuses these representations using a cross-attention mechanism in which the attention logits are augmented with a bias derived from LJ. The resulting fused embedding is subsequently used to condition a denoising diffusion probabilistic model (DDPM). During the diffusion process, a random latent vector is progressively refined into a conformationally aware embedding that captures the system’s structural flexibility. Finally, an SE(3)-Equivariant decoder translates this refined embedding into the final 3D atomic coordinates of the bound complex. The detailed architecture of the decoder can be found in SI Appendix 1.C. The entire framework is trained in an end-to-end manner under a multi-objective loss function that simultaneously optimizes for structural accuracy, diffusion denoising performance, and fundamental physical and geometric constraints, ensuring the generation of biophysically realistic structures.

### Cooperative Representation Encoding Module

As shown in Figure 1, CoupleDiff employs a representation encoding module composed of two main components: cooperative scalar-vector representation and protein embedding discretization to represent both the ligand and protein molecules. Both protein and ligand are represented as graphs. We utilize hierarchical encoding for the protein by dividing it into the backbone *G*^*b*^ = ( *V*^*b*^, *ℰ*^*b*^) and the side chain *G*^*s*^ = ( *V*^*s*^, *ℰ*^*s*^). All nodes and edges are parameterized by scalar and vector features (as shown in SI Appendix 1.B).

### Protein Embedding Discretization

After obtaining the enhanced embedding via scalar-vector cooperation, the protein representation is discretized using the pre-trained FoldToken4 codebook (42). This discrete encoding has been shown to more effectively capture recurring structural motifs, reduce representational variance, and minimize redundancy in the learned embeddings. Using the inherent regularity of the backbone structures compared to the conformational diversity of the side chains, we map the continuous geometric embedding *e*^*b*^ into a discrete semantic space through vector quantization. Specifically, the high-dimensional continuous representation *e*^*b*^ is compressed into a discrete semantic token *e*^*t*^:

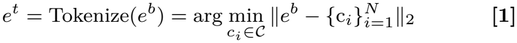

by minimizing the Euclidean reconstruction error between the backbone embedding and the codebook vectors *C* = {*c*_1_, *c*_2_, … , *c*_*N*_}. This discretization mechanism achieves dimension-reduced encoding of 3D structures while preserving backbone topological features, significantly enhancing storage efficiency and computational tractability of protein representations.

### Non-Covalent Interaction Module

In this section, we adopt the non-covalent interaction module to generate an adaptive joint embedding for the protein–ligand complex.

To capture the adaptive nature of protein–ligand interactions, we incorporate docking-derived interaction priors into the attention computation. Specifically, we extend the conventional cross-attention formulation by integrating a physics-constrained interaction matrix that reflects atomic-level docking scores, as shown in Equation 2:

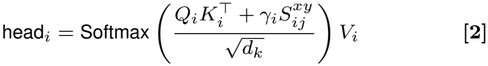

Here, 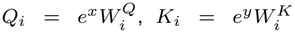, and 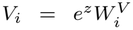, where *x, y, z* ∈ {*b, s, l*} denote the backbone (*b*), side-chain (*s*), and ligand (*l*) embeddings respectively, depending on the directional pair under consideration. Let 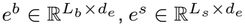 , and 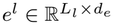 be the corresponding input features, with 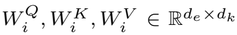 denoting learnable projection matrices. *L*_*b*_ represents the number of nodes in the protein backbone. Each node in the backbone graph corresponds to a single residue. *L*_*s*_ represents the number of nodes in the protein side-chain. In our atom-level graph for the side-chain, each node corresponds to a single heavy atom. *d*_*e*_ represents the dimensionality of the node feature embeddings. It is a hyperparameter that defines the length of the feature vector used to represent each node, whether it is a residue in the backbone or an atom in the side-chains or the ligand. The learnable scalar *γ*_*i*_ controls the influence of the physical prior on the final attention distribution.

The interaction matrix 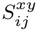 is computed by using LJ potentials of protein– ligand conformation modeled by AutoDock tool. For each directional pair *x, y* ∈ {*b, s, l*}, we define

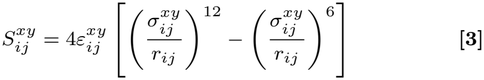

where *r*_*ij*_ = ∥*p*_*i*_ ™ *p*_*j*_ ∥ is the Euclidean distance. 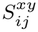serves as an interaction-aware prior that modulates the raw attention logits. By adding 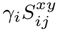 to 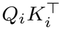, the model is guided to emphasize spatially and chemicall plausible interaction sites, promoting more meaningful attention across protein–ligand pairs. And the LJ parameters are computed via Lorentz-Berthelot rules:

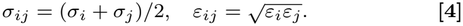

where *σ*_*i*_ and *σ*_*j*_ are the LJ size (collision-diameter) parameters of atom *i* and atom *j. ε*_*i*_ and *ε*_*j*_ are the LJ energy-well depth parameters of atom *i* and *j*.

To account for the full interaction landscape, we compute attention over six directional combinations: *b* → *s, s* → *b, b* → *l, l* → *b, s* → *l*, and *l* → *s*. Each head output has shape 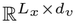 . Concatenating all six directional outputs along the feature dimension yields a tensor of shape 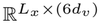, which is then passed through a trainable MLP to project it back into 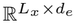 to produce a compact fused embedding *e*^*f*^ . By integrating these interaction-aware priors, the model generates adaptive, ligand-specific representations that reflect diverse structural or functional responses across protein environments.

### Physics-Guided Diffusion Module

To explicitly model the adaptive and physically constrained nature of protein–ligand binding, we propose a physics-guided diffusion module that integrates conditional latent diffusion with geometry- and force-aware supervision. This design enables the model to generate conformation-aware embeddings that are not only expressive but also biophysically plausible.

### Conditional Diffusion in Latent Interaction Space

Given the fused protein–ligand embedding *e*^*f*^ , we employ a conditional DDPM to generate a refined embedding *e*_refined_ that captures structural adaptability. The diffusion process operates on a latent variable *z*, allowing flexible modeling of interactions while decoupling generative uncertainty from coordinate space.

The forward diffusion process *q* progressively perturbs the ground-truth embedding *e*_*gt*_ with Gaussian noise over *T* timesteps, starting from *z*_0_ = *e*_*gt*_:

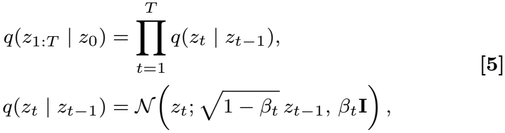

where 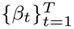 is a predefined noise schedule.

The reverse denoising process is parameterized by a neural network *p*_*θ*_, which iteratively reconstructs clean embeddings from noisy samples while being explicitly conditioned on the fused interaction context *e*^*f*^

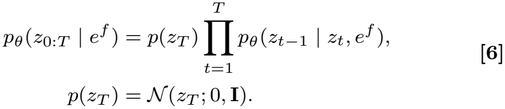

During inference, we initialize from isotropic Gaussian noise *z*_*T*_ ∼ *N* (0, **I**) and apply the learned denoising transitions from *t* = *T* to 1, yielding the refined embedding *e*_refined_ = *z*_0_. Conditioning on *e*^*f*^ at each step ensures that the diffusion trajectory remains anchored to the specific protein–ligand pair, enabling protein-dependent conformational refinement. **Force- and Geometry-Guided Supervision**. While diffusion provides a powerful mechanism for modeling uncertainty and flexibility, unconstrained denoising in latent space may lead to physically implausible structures when decoded. To address this, we explicitly guide the diffusion process using force- and geometry-aware losses that regularize both the embedding evolution and the decoded atomic structures.

Specifically, we introduce a physics and geometry-guided loss ℒ_phy-geo_ composed of seven terms that encode fundamental molecular constraints, including bond lengths, bond angles, dihedral angles, vdW interactions, electrostatics, hydrogen bonding, and *π*–*π* stacking interactions:

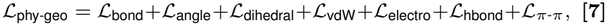

with detailed formulations provided in SI Appendix 1.D. These terms collectively enforce chemically feasible local geometry and physically meaningful non-covalent interactions.

In addition to interaction-level regularization, we supervise the decoded all-atom complex structure by minimizing the mean squared deviation between predicted atomic coordinates 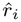 and ground-truth positions *r*_*i*_:

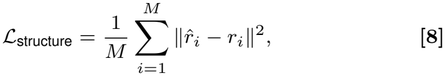

where *M* denotes the number of atoms in the complex.

Finally, to train the diffusion model itself, we adopt the standard noise prediction objective, which encourages accurate denoising of corrupted embeddings at each timestep *t*:

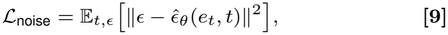

The overall training objective integrates diffusion consistency, structural accuracy, and physical plausibility into a unified framework:

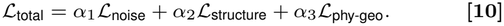

where *α*_1_, *α*_2_, and *α*_3_ are adjusted weights that balance generative fidelity and physical realism throughout training (see SI Appendix 3.G).

By embedding physical and geometric constraints into the diffusion-based refinement process, our method controls the stochastic denoising trajectory toward energetically favorable and structurally feasible regions of the interaction, enabling accurate modeling of adaptive protein–ligand binding. Critically, these physical and geometric terms are not confined to training: during inference they act as explicit guidance on the reverse-diffusion trajectory, steering each denoising step toward physically compatible configurations.

## Data, Materials, and Software Availability

The source code for CoupleDiff is available at https://github.com/DiyaZhang6/CoupleDiff. Additional details are provided in *SI Appendix*.

## ACKNOWLEDGMENTS

This work was supported by the National Natural Science Foundation of China (62322215, 62532017, 62602557, 62402488, 62372279), the Natural Science Foundation of Hunan Province (2025JJ60387), and the Natural Science Foundation of Shandong Province (ZR2025QB62, ZR2023MF119). This study was also supported in part by the High-Performance Computing Center of Central South University.

