## Supplementary Information for "Physics-Guided Generative Modeling Reveals that Steric Conflict Resolution Drives Ligand Specificity and Induced-Fit Adaptation"

#### This PDF file includes:

Supporting text
Figs. S1 to S6
Tables S1 to S25
SI References

### Supporting Information Text

### 1. CoupleDiff Architecture

**A. Graph Feature Construction.** Both target and drug are represented as graphs. We utilize hierarchical encoding for the target by dividing it into the backbone $\mathcal{G}^b = (\mathcal{V}^b, \mathcal{E}^b)$ and the side chain $\mathcal{G}^s = (\mathcal{V}^s, \mathcal{E}^s)$. All nodes and edges are jointly parameterized by scalar and vector features. For target, the backbone graph encodes the global structural scaffold, preserving its rigid tertiary folding through continuous peptide bond connectivity. For each node, scalar features include amino acid type, residue ID from the sequence, hydrophobicity and secondary structure, while vector features incorporate geometric information including Cartesian coordinates of the $C_\alpha$ atom, two dihedral angles and four direction vectors. Foe each edge, scalar edge features include Euclidean distance between two adjacent residues and bond type. Vector edge features include the direction vector of two adjacent residues. Meanwhile, the side-chain graph $\mathcal{G}^s = (\mathcal{V}^s, \mathcal{E}^s)$ of target captures localized chemical environments at binding sites, representing rotatable bonds and functional group orientations critical for molecular recognition. Scalar features include atom type, atom number, B-factor, functional groups type, implicit valence electrons, aromaticity, electrostatic charge and hybridization state, while vector features include the atom coordinates and five dihedral angles, which are represented by their sine and cosine values. Scalar edge features are same as the backbone. The atomic graph $\mathcal{G}^s = (\mathcal{V}^s, \mathcal{E}^s)$ explicitly models conformationally flexible side-chain atoms (e.g., $\chi$ -angle rotations in tyrosine residues), focusing on their spatial rearrangements during drug binding (1). Additionally, the atomic graph $\mathcal{G}^d$ is also represented by physical and chemical properties and geometric features, such as element type, partial charge, hybridization state, chirality, aromaticity, atomic mass.

**B. Cooperative Scalar-Vector Representation Details.** To overcome the decoupling of scalar and vector channels in the original SE(3)-Transformer (2), we devise a bidirectional update mechanism that couples geometric features, represented as vectors, with semantic features, represented as scalars, as illustrated in Algorithm 1. At each iteration, the passing of the node-node and edge-edge message is enhanced by gated updates: for each node $v_i$, we first compute the displacement vector between neighboring nodes and transform it into spherical coordinates $(\theta_{ij}, \phi_{ij})$ to provide a rotation-aware geometric basis. We then compute directional messages $\mathbf{h}_{ij}$ from neighbors $v_j$ that fuse its scalar descriptor, edge scalars and a learned encoding of the edge vector; attention weights $\alpha_{ij}$ then modulate their aggregation into a residual message $\mathbf{m}_i$. The node scalar feature is updated by a multi-layer perceptron (MLP), while its vector feature undergoes an SE(3)-equivariant correction via another MLP. Edge features are likewise refined: spherical harmonics-based geometric embeddings $\mathbf{h}_{ij}^{\text{geometric}}$ augment edge scalars through an edge-MLP, and the edge vector is shifted by a learned function of its incident node vectors.

Finally, an interaction module executes a scalar-vector inner product at each node to produce a cross-modal message $\mathbf{h}_{\text{interact}}$, which is fed back to both channels, adding squared norms of the other channel as residual boosts, to enable true bidirectional flow. We apply this cooperative encoding to the ligand graph $\mathcal{G}^l$, as well as the protein backbone $\mathcal{G}^b$ and side-chain $\mathcal{G}^s$ graphs. After $T$ iterations, spatially aggregated features (via global average pooling over nodes and edges) are passed through task-specific MLP to yield three embeddings: the backbone embedding $e^b$ (global fold topology), the side-chain embedding $e^s$ (local conformational variability), and the ligand embedding $e^l$.

**C. Decoder Architecture.** The decoder $f_{\text{dec}}$ is responsible for transforming the final diffusion-denoised fused representation $e_f^0 \in \mathbb{R}^{N\times d}$—which jointly encodes physics-aware target-drug interaction semantics—into a physically plausible 3D structure $\hat{C} = \{\hat{r}_i \in \mathbb{R}^3\}_{i=1}^N$, where $N$ denotes the number of nodes (residues and atoms).

The decoder is implemented as an $L$-layer SE(3)-Equivariant graph transformer, where each node $i$ is initialized with the latent feature $h_i^{(0)} = e_f^{0,i}$ from diffusion and an initial coordinate $r_i^{(0)}$ obtained via a pre-docking strategy:

$$h_i^{(0)} = W^{(0)} e_f^{0,i} + b^{(0)}, \quad r_i^{(0)} = r_i^{\text{init}} \qquad [1]$$

At each layer $\ell = 1, \ldots, L$, the decoder performs the following operations.

***C.1. Equivariant Attention.*** At each layer $\ell$, the decoder refines node-level representations through a self-attention mechanism that integrates contextual information across the target–drug interaction embedding. Specifically, each node $i$ with scalar feature $h_i^{(\ell-1)} \in \mathbb{R}^d$ computes attention weights over its neighbors by projecting its embedding into a query vector:

$$Q_i^{(\ell)} = h_i^{(\ell-1)} W_Q^{(\ell)}, \qquad [2]$$

where $W_Q^{(\ell)} \in \mathbb{R}^{d\times d}$ is a learnable projection matrix at layer $\ell$.

Similarly, all other nodes $j$ are projected into key and value vectors:

$$K_j^{(\ell)} = h_j^{(\ell-1)} W_K^{(\ell)}, \quad V_j^{(\ell)} = h_j^{(\ell-1)} W_V^{(\ell)} \qquad [3]$$

where $W_K^{(\ell)}, W_V^{(\ell)} \in \mathbb{R}^{d\times d}$ are the key and value projection matrices.

The attention weight between node $i$ and node $j$ is computed via scaled dot-product attention:

$$a_{ij}^{(\ell)} = \frac{\exp\left(Q_i^{(\ell)} \cdot K_j^{(\ell)} / \sqrt{d}\right)}{\sum_{k=1}^N \exp\left(Q_i^{(\ell)} \cdot K_k^{(\ell)} / \sqrt{d}\right)}, \qquad [4]$$

**Algorithm 1** Scalar-Vector Feature Interaction with SE(3)-Equivariant Update

**Input:**
Node scalar features $\mathbf{f}_{\text{scalar}}(v_i)$, vector features $\mathbf{f}_{\text{vector}}(v_i)$ for all $v_i \in \mathcal{V}$
Edge scalar features $\mathbf{f}_{\text{scalar}}(e_{ij})$, vector features $\mathbf{f}_{\text{vector}}(e_{ij})$ for all $e_{ij} \in \mathcal{E}$
**Output:** Updated $\mathbf{f}_{\text{scalar}}(v_i)$, $\mathbf{f}_{\text{vector}}(v_i)$, $\mathbf{f}_{\text{scalar}}(e_{ij})$, $\mathbf{f}_{\text{vector}}(e_{ij})$
Initialize iteration $t \leftarrow 0$, maximum iterations $T$
**repeat**
  **for** each node $v_i \in \mathcal{V}$ **do**
    **for** each neighbor $v_j \in \mathcal{N}(v_i)$ **do**
      $\mathbf{r}_{ij} \leftarrow \mathbf{f}_{\text{vector}}(v_j) - \mathbf{f}_{\text{vector}}(v_i)$
      $\theta_{ij}, \phi_{ij} \leftarrow \textsc{SphericalCoordinates}(\mathbf{r}_{ij})$
      $\mathbf{h}_{ij} \leftarrow \textsc{Concat}\big(\mathbf{f}_{\text{scalar}}(v_i), \mathbf{f}_{\text{scalar}}(e_{ij}), \text{MLP}(\mathbf{f}_{\text{vector}}(e_{ij}))\big)$
      $\alpha_{ij} \leftarrow \textsc{Softmax}\left(\mathbf{W}_1^\top \mathbf{h}_{ij} + \mathbf{b}_1\right)$
    $\mathbf{m}_i \leftarrow \sum_{j\in\mathcal{N}(v_i)} \alpha_{ij} \cdot \mathbf{h}_{ij}$
    $\mathbf{f}_{\text{scalar}}(v_i) \leftarrow \text{MLP}_{\text{node}}\left(\mathbf{f}_{\text{scalar}}(v_i), \mathbf{m}_i\right)$
    $\mathbf{f}_{\text{vector}}(v_i) \leftarrow \mathbf{f}_{\text{vector}}(v_i) + \text{MLP}_{\text{vec}}(\mathbf{m}_i)$
  **for** each edge $e_{ij} \in \mathcal{E}$ **do**
    $\mathbf{h}_{ij}^{\text{geometric}} \leftarrow \sum_{l=0}^{L} \sum_{m=-l}^{l} w_{lm} N_{lm} P_l^m(\cos\theta_{ij}) e^{im\phi_{ij}}$
    $\mathbf{f}_{\text{scalar}}(e_{ij}) \leftarrow \text{MLP}_{\text{edge}}\left(\mathbf{f}_{\text{scalar}}(e_{ij}), \mathbf{h}_{ij}^{\text{geometric}}\right)$
    $\mathbf{f}_{\text{vector}}(e_{ij}) \leftarrow \mathbf{f}_{\text{vector}}(e_{ij}) + \text{MLP}_{\text{vec-edge}}(\mathbf{f}_{\text{vector}}(v_i), \mathbf{f}_{\text{vector}}(v_j))$
**Interaction Module:**
  **for** each node $v_i \in \mathcal{V}$ **do**
    $\mathbf{h}_{\text{interact}} \leftarrow \text{MLP}(\mathbf{f}_{\text{scalar}}(v_i)) \cdot \mathbf{f}_{\text{vector}}(v_i)$
    $\mathbf{f}_{\text{scalar}}(v_i) \leftarrow \mathbf{f}_{\text{scalar}}(v_i) + \|\mathbf{f}_{\text{vector}}(v_i)\|_2^2$
    $\mathbf{f}_{\text{vector}}(v_i) \leftarrow \mathbf{f}_{\text{vector}}(v_i) + \mathbf{h}_{\text{interact}} \cdot \|\mathbf{f}_{\text{scalar}}(v_i)\|_2$
  $t \leftarrow t + 1$
**until** $t \geq T$

where $d$ is the hidden dimension and $a_{ij}^{(\ell)}$ defines the normalized attention score assigned to node $j$ when updating node $i$.

Using these weights, node $i$ aggregates information from all neighbors:

$$\tilde{h}_i^{(\ell)} = \sum_{j=1}^{N} a_{ij}^{(\ell)} V_j^{(\ell)}, \quad [5]$$

where $\tilde{h}_i^{(\ell)}$ represents the aggregated message from the neighborhood of $i$.

Finally, the new node representation is updated using a residual connection followed by layer normalization:

$$h_i^{(\ell)} = \text{LayerNorm}\left(h_i^{(\ell-1)} + \tilde{h}_i^{(\ell)}\right), \quad [6]$$

where $h_i^{(\ell)}$ is the refined embedding at layer $\ell$.

***C.2. Message Passing.*** To incorporate distance-aware relational information into node-level reasoning, we employ a message passing mechanism that encodes pairwise Euclidean distances into high-dimensional edge features. At each layer $\ell$, for each node pair $(i, j)$, we compute their spatial distance based on current coordinates:

$$d_{ij}^{(\ell-1)} = \|r_i^{(\ell-1)} - r_j^{(\ell-1)}\|_2, \quad [7]$$

where $r_i^{(\ell-1)} \in \mathbb{R}^3$ is the coordinate of node $i$ at layer $\ell-1$, and $d_{ij}^{(\ell-1)}$ denotes the Euclidean distance between nodes $i$ and $j$.

The scalar distance $d_{ij}^{(\ell-1)}$ is then passed through a radial basis function expansion to obtain a smooth, high-dimensional representation:

$$\phi_k(d_{ij}) = \exp\left(-\beta_k (d_{ij} - \mu_k)^2\right), \quad k = 1, \ldots, K, \quad [8]$$

where $\mu_k$ and $\beta_k$ are the center and bandwidth of the $k$-th Gaussian kernel, respectively. The output $\{\phi_k(d_{ij})\}_{k=1}^K$ is a $K$-dimensional vector encoding the relative spatial proximity between nodes $i$ and $j$.

Finally, we compute the edge-wise message $m_{ij}^{(\ell)}$ by integrating both node-level semantic information and the geometric encoding:

$$m_{ij}^{(\ell)} = \phi_m\left(h_i^{(\ell)},\ h_j^{(\ell)},\ \{\phi_k(d_{ij})\}_{k=1}^K\right), \quad [9]$$

where $h_i^{(\ell)} \in \mathbb{R}^d$ and $h_j^{(\ell)} \in \mathbb{R}^d$ are the current scalar embeddings of nodes $i$ and $j$, and $\phi_m(\cdot)$ denotes a learnable multi-layer perceptron (MLP) that fuses semantic and geometric inputs into a unified message representation.

***C.3. Coordinate Update.*** To refine the atomic coordinates in an SE(3)-Equivariant manner, we apply a direction-aware coordinate update based on the message $m_{ij}^{(\ell)}$ computed in C.2. For each node pair $(i, j)$, a scalar weight $w_{ij}^{(\ell)}$ is first predicted via a learnable neural function:

$$w_{ij}^{(\ell)} = \psi^{(\ell)}(m_{ij}^{(\ell)}), \quad [10]$$

where $\psi^{(\ell)}(\cdot)$ is an MLP that maps the message $m_{ij}^{(\ell)}$ into a scalar attention score that modulates the geometric influence of node $j$ on node $i$.

Next, the coordinate displacement for node $i$ is computed by aggregating normalized directional vectors weighted by $w_{ij}^{(\ell)}$:

$$\Delta r_i^{(\ell)} = \sum_j w_{ij}^{(\ell)} \cdot \frac{r_i^{(\ell-1)} - r_j^{(\ell-1)}}{d_{ij}}, \quad [11]$$

where $r_i^{(\ell-1)} \in \mathbb{R}^3$ and $r_j^{(\ell-1)} \in \mathbb{R}^3$ are the coordinates of nodes $i$ and $j$ at layer $\ell - 1$, and $d_{ij}$ is their Euclidean distance as previously defined. The term $(r_i - r_j)/d_{ij}$ is the unit direction vector pointing from $j$ to $i$, ensuring the update direction is geometrically meaningful and rotation-equivariant.

Finally, the coordinates of node $i$ are updated via residual addition:

$$r_i^{(\ell)} = r_i^{(\ell-1)} + \Delta r_i^{(\ell)}, \quad [12]$$

producing refined coordinates $r_i^{(\ell)}$ at layer $\ell$. The update formulation allows gradual correction of initial positions while preserving SE(3) equivariance, as the entire update depends on relative positions and scalar weights.

***C.4. Final Prediction.*** After $L$ layers of equivariant updates, each node $i$ is associated with a refined coordinate $r_i^{(L)} \in \mathbb{R}^3$. To ensure the final predicted structure is invariant to global translations, we subtract the centroid of all node positions before output:

$$\hat{r}_i = r_i^{(L)} - \frac{1}{N}\sum_{j=1}^{N} r_j^{(L)}, \qquad \hat{C} = \{\hat{r}_i\}_{i=1}^{N} \quad [13]$$

where $N$ denotes the number of nodes, and $\hat{r}_i \in \mathbb{R}^3$ is the final translation-normalized coordinate of node $i$. The term $\frac{1}{N}\sum_{j=1}^{N} r_j^{(L)}$ represents the centroid of the entire structure at the final layer, and its subtraction centers the predicted coordinates around the origin.

This normalization step guarantees that the predicted structure $\hat{C}$ remains invariant under global translations of the input coordinate frame, preserving the SE(3)-Equivariant property of the overall architecture.

**D. Multi-Objective Loss and Physical Geometry Functions.** The coordinate predictions $\hat{C}$ are supervised under a multi-objective loss function that enforces physical plausibility.

Bond lengths are constrained by introducing a penalty term that discourages deviations from their reference covalent bond distances, which are typically in the range of 1.0-2.5 Å:

$$\mathcal{L}_{\text{bond}} = \sum_{(i,j)\in\mathcal{B}} \left( \frac{\|\mathbf{r}_i - \mathbf{r}_j\|_2 - d_{ij}^{\text{ref}}}{d_{ij}^{\text{ref}}} \right)^2 \quad [14]$$

where $\mathcal{B}$ denotes the set of bonded atom pairs, and $d_{ij}^{\text{ref}}$ are experimentally derived bond lengths.

Bond angle consistency maintains directional geometry by constraining angles (102°–120°) between consecutive bonds:

$$\mathcal{L}_{\text{angle}} = \sum_{(i,j,k)\in\mathcal{A}} \left( \cos^{-1}\left( \frac{(\mathbf{r}_i - \mathbf{r}_j)\cdot(\mathbf{r}_k - \mathbf{r}_j)}{\|\mathbf{r}_i - \mathbf{r}_j\|_2 \|\mathbf{r}_k - \mathbf{r}_j\|_2} \right) - \theta_{ijk}^{\text{ref}} \right)^2 \quad [15]$$

where $\mathcal{A}$ represents valid bond angle triplets.

To preserve the higher-order structural stability of the molecule, dihedral angle consistency constrains the torsional rotation between four sequentially bonded atoms, capturing the flexibility of the target and drug conformation:

$$\mathcal{L}_{\text{dihedral}} = \sum (i,j,k,l) \in \mathcal{D} \left( \phi_{ijkl}^{\text{pred}} - \phi_{ijkl}^{\text{true}} \right)^2 \quad [16]$$

where $\mathcal{D}$ denotes the all relevant dihedral angle quadruplets, where each quadruplet $(i, j, k, l)$ corresponds to four sequentially bonded atoms defining a torsional degree of freedom. The term $\phi^{\text{pred}} ijkl$ represents the predicted dihedral angle for the quadruplet formed by atoms $i$, $j$, $k$, and $l$, while $\phi^{\text{true}} ijkl$ corresponds to the reference torsional angle for the same set of atoms.

Steric exclusion enforcement ensures that no two atoms occupy the same physical space by applying vdW radius-based thresholds:

$$\mathcal{L}_{\text{vdW}} = \sum_{i<j} \text{ReLU}\left( r_{ij}^{vdW} - \| \mathrm{r}_i - \mathrm{r}_j \|_2 \right)^2 \quad [17]$$

where $r_{ij}^{vdW}$ is the sum of atomic vdW radii.

To account for long-range Coulombic effects in the target-drug complex, we introduce a dipole–dipole interaction term based on the leading order of multipole expansion. This term enforces electrostatic plausibility by penalizing deviations from physically consistent dipole alignments:

$$\mathcal{L}_{\text{electro}} = \sum_{(i,j)\in\mathcal{P}} \left( \frac{\mathbf{p}_i \cdot \mathbf{p}_j - 3\,(\mathbf{p}_i \cdot \hat{\mathbf{r}}_{ij})\,(\mathbf{p}_j \cdot \hat{\mathbf{r}}_{ij})}{\|\mathbf{r}_i - \mathbf{r}_j\|_2^3} \right)^2 \tag{18}$$

where $\mathcal{P}$ denotes all relevant dipole–dipole pairs, $\mathbf{p}_i$ and $\mathbf{p}_j$ are the atomic dipole moments, and $\hat{\mathbf{r}}_{ij}$ is the unit vector from atom $i$ to atom $j$.

Hydrogen bonding geometry is regularized to reflect canonical interaction patterns, enforcing appropriate donor–acceptor distances and angles between participating atoms:

$$\begin{aligned}\mathcal{L}_{\text{hbond}} = \sum_{(d,h,a)\in\mathcal{H}} \Big[ &\text{ReLU}\big(|\boldsymbol{r}_d - \boldsymbol{r}_a| - 3.2\big)^2 \\ &+ \text{ReLU}\big(2.5 - |\boldsymbol{r}_d - \boldsymbol{r}_a|\big)^2 \\ &+ \text{ReLU}\big(120 - \theta\big)^2 + \text{ReLU}\big(\theta - 180\big)^2 \Big]\end{aligned} \tag{19}$$

where $\mathcal{H}$ is the set of donor-hydrogen-acceptor triplets, $\boldsymbol{r}_d$ and $\boldsymbol{r}_a$ are donor and acceptor coordinates, and $\theta$ is the donor-hydrogen-acceptor angle.

We define a $\pi - \pi$ interaction loss $\mathcal{L}_{\pi-\pi}$ that penalizes deviations from ideal aromatic stacking geometries and alignment:

$$\begin{aligned}\mathcal{L}_{\pi\pi\text{-geo}} = \sum_{(i,j)\in\pi\pi \text{ pairs}} \Bigg[ &\text{ReLU}(d_{ij} - 5.0)^2 + \text{ReLU}(3.5 - d_{ij})^2 \\ &+ \text{ReLU}(|\theta_{ij}| - 30^\circ)^2 + \text{ReLU}(|\Delta x_{ij}| - 3.5)^2 \Bigg]\end{aligned} \tag{20}$$

$$\mathcal{L}_{\pi-\pi} = \mathcal{L}_{\pi\pi\text{-geo}} + \mathcal{L}_{\pi\pi\text{-align}} \tag{21}$$

where $(i, j)$ are pairs of aromatic rings involved in $\pi$-$\pi$ interactions, $d_{ij}$ is the distance between centroids, $\theta_{ij}$ is the inter-centroid angle, $\Delta x_{ij}$ is the lateral displacement, and $\boldsymbol{n}_i, \boldsymbol{n}_j$ are normal vectors of the aromatic planes.

**E. Overall Inference Pipeline.** The overall inference pipeline of the proposed framework, which takes a target protein and a drug molecule as input and outputs a refined three-dimensional binding pose, is summarized in Algorithm 2.

**Algorithm 2** Structure Refinement via Diffusion and Geometric Constraints

**Input:** Target structure $P$, drug SMILES $S$
**Output:** Predicted binding pose $\hat{C}$ (3D coordinates)
Convert $S$ to 3D drug structure $D$ via RDKit
Generate initial pose $C_0$ by docking $P$ and $D$ with AutoDock
Build backbone graph $\mathcal{G}^b$ and side-chain graph $\mathcal{G}^s$ from $P$
Build drug graph $\mathcal{G}^D$ from $D$
$e^b \leftarrow \text{Encoder}_{\text{backbone}}(\mathcal{G}^b)$
$e^d \leftarrow \text{Encoder}_{\text{drug}}(\mathcal{G}^D)$
$e^s \leftarrow \text{Encoder}_{\text{side-chain}}(\mathcal{G}^s)$
$e^t \leftarrow \text{Tokenize}(e^b)$
$e^f \leftarrow \text{PhysicsAttention}(e^t, e^d)$
$e_f \leftarrow \text{Linear}(e^f)$
Sample noise vector $z_T \sim \mathcal{N}(0, \mathbf{I})$
**for** $t = T$ down to 1 **do**
    $\hat{\boldsymbol{\epsilon}}_\theta \leftarrow \text{DenoisingNet}_\theta(z_t, t, e_f)$
    $z_{t-1} \leftarrow \frac{1}{\sqrt{\alpha_t}} \left( z_t - \frac{1-\alpha_t}{\sqrt{1-\bar{\alpha}_t}} \hat{\boldsymbol{\epsilon}}_\theta \right)$
$e_{\text{refined}} \leftarrow z_0$
$\hat{C} \leftarrow \text{Decoder}(e_{\text{refined}})$

Given a target structure $P$ and a drug specified by its simplified molecular input line entry system (SMILES) string $S$, we first construct an initial drug conformation, followed by a docking step to generate a coarse binding pose. This initialization provides a physically reasonable starting point for subsequent refinement.

To encode structural information at multiple resolutions, the target is represented as two complementary graphs corresponding to backbone and side-chain structures, while the drug is modeled as a molecular graph. These components are processed by encoders to obtain latent representations capturing both global topology and local chemical environments. The backbone representation is further tokenized and fused with ligand features via a physics-aware attention mechanism, enabling effective cross-modal interaction.

Conditioned on the fused representation, we perform a diffusion-based refinement process in latent space. Starting from Gaussian noise, the model iteratively denoises the latent variable using a parameterized network, progressively reconstructing a representation that is consistent with both geometric constraints and learned interaction patterns.

Finally, the refined latent representation is decoded into three-dimensional coordinates, yielding the predicted binding pose.

### 2. Details of Datasets

We have performed a detailed analysis of our full learnable dataset (Table S1), confirm that the dataset is sufficiently large and diverse to support the learning of generalized principles of target-drug interaction.

**Table S1. Statistics of our curated dataset.**

| Category | Metric | Value |
|---|---|---|
| Overall Dataset Size | Total Complexes | 21,762 |
| | Unique Protein Targets | 3,673 |
| Ligand Diversity | Mean Unique Ligands per Target | 6.57 |
| | Median Unique Ligands per Target | 3.0 |
| | Targets with > 5 Unique Ligands | 917 |
| | Targets with > 10 Unique Ligands | 444 |
| | Targets with > 20 Unique Ligands | 200 |
| Structural Diversity | Targets with > 5 Co-crystal Structures | 609 |
| | Targets with > 10 Co-crystal Structures | 318 |
| | Targets with > 20 Co-crystal Structures | 148 |
| Protein Family Diversity | HYDROLASE | 16.6% |
| | TRANSFERASE | 16.0% |
| | HYDROLASE/INHIBITOR | 10.3% |
| | TRANSFERASE/INHIBITOR | 10.1% |
| | TRANSCRIPTION | 3.6% |

### 3. Detailed Experimental Results

**A. Results of Redocking Task.** We evaluate the performance of various computational methods on the redocking task, which involves reconstructing the three-dimensional binding conformations of target proteins and their corresponding drugs starting from experimentally determined structures, in order to assess each method's accuracy and reliability in reproducing native complex geometries.

On our curated test set (Table S3), CoupleDiff achieves the best performance across all evaluation metrics, including overall, side-chain, and ligand root mean square deviation (RMSD), as well as success rates under the 2 Å threshold. In particular, it delivers consistent improvements across both RMSD-based accuracy and success rate metrics, indicating not only reduced structural deviation but also a higher likelihood of generating near-native binding conformations. Compared with existing docking approaches and recent learning-based methods, CoupleDiff shows advantages in drug reconstruction while maintaining strong performance in target-side structural modeling. This improvement across different components of the complex suggests that the model can effectively coordinate drug placement with the surrounding target environment.

On the Astex benchmark (Table S5), CoupleDiff further demonstrates strong generalization ability. It outperforms all competing methods with a clear margin in both RMSD and success rate metrics, suggesting that the model is robust to diverse binding environments and molecular compositions. Compared with traditional docking approaches and recent learning-based methods, CoupleDiff achieves more balanced improvements across backbone, side-chain, and drug reconstruction.

For the more challenging APObind setting (Table S7), where targets are provided in their apo forms, all methods experience noticeable performance degradation. Nevertheless, CoupleDiff maintains the best overall performance, achieving the lowest RMSD and highest success rates. This result highlights its ability to handle structural flexibility and infer induced-fit effects, which are critical for realistic drug discovery scenarios.

Finally, on the ESMFold-predicted structures (Table S9), which introduce additional uncertainty from predicted protein backbones, CoupleDiff remains consistently competitive and achieves the best overall performance. Despite the increased noise

**Table S3. Performance of different methods in reconstructing complex structures on our curated test set, evaluated by RMSD (Å) and success rate.**

| Method | RMSD (↓) | | | Success Rate (↑) | | |
|---|---|---|---|---|---|---|
| | Overall | Side-chain | Ligand | Overall RMSD $< 2$ Å | sc-RMSD $< 2$ Å | L-RMSD $< 2$ Å |
| Glide (3) | 11.20±4.92 | 9.37±4.32 | 15.01±4.98 | 18.79% | 19.73% | 10.31% |
| RosettaLigand (4) | 9.83±4.01 | 9.19±4.13 | 19.32±4.84 | 20.97% | 21.02% | 8.33% |
| AutoDock (5) | 8.60±4.73 | 7.05±4.01 | 13.29±4.56 | 21.06% | 22.80% | 12.56% |
| TankBind (6) | 5.36±3.14 | 4.48±3.25 | 14.66±5.08 | 23.51% | 29.10% | 11.34% |
| DiffDock (7) | 3.77±2.85 | 2.53±1.40 | 9.73±4.10 | 35.71% | 57.50% | 18.00% |
| DiffDock-Pocket (8) | 3.19±1.40 | 1.27±1.56 | 6.24±4.22 | 39.10% | 67.20% | 25.65% |
| DynamicBind (9) | 3.01±1.88 | 1.36±0.67 | 3.25±2.67 | 38.72% | 70.10% | 51.77% |
| DiffBindFR (1) | 2.98±1.38 | 0.61±0.38 | 2.73±1.30 | 50.98% | 55.30% | 57.22% |
| SurfDock (10) | 2.55±2.35 | 0.55±1.37 | 4.59±2.46 | 58.23% | 70.00% | 46.16% |
| PackDock (11) | 2.52±1.44 | 0.52±0.37 | 2.80±1.42 | 56.54% | 68.10% | 57.89% |
| FlexPose (12) | 2.44±2.83 | 0.44±0.34 | 3.48±2.20 | 60.17% | 70.20% | 30.24% |
| **CoupleDiff** | **2.01±1.85** | **0.29±1.04** | **2.17±1.45** | **65.30%** | **70.50%** | **61.60%** |

(↑) / (↓) denotes higher / lower is better; Top 1 and Top 2 are highlighted with **bold** and underlined, respectively.

**Table S5. Performance of different methods in reconstructing complex structures on the Astex set, evaluated by RMSD (Å) and success rate.**

| Method | RMSD (↓) | | | Success Rate (↑) | | |
|---|---|---|---|---|---|---|
| | Overall | Side-chain | Ligand | Overall RMSD $< 2$ Å | sc-RMSD $< 2$ Å | L-RMSD $< 2$ Å |
| Glide (3) | 2.63±1.46 | 0.44±1.03 | 3.28±1.94 | 55.01% | 57.45% | 49.28% |
| RosettaLigand (4) | 2.74±1.53 | 0.45±0.19 | 4.26±1.43 | 56.29% | 58.37% | 45.20% |
| AutoDock (5) | 2.54±0.99 | 0.72±0.18 | 5.90±1.67 | 50.62% | 54.29% | 22.38% |
| TankBind (6) | 7.36±4.82 | 6.25±2.75 | 12.52±3.63 | 22.61% | 25.07% | 16.46% |
| DiffDock (7) | 3.38±1.20 | 3.21±1.66 | 8.20±2.22 | 37.93% | 38.27% | 20.19% |
| DiffDock-Pocket (8) | 3.19±0.38 | 3.08±1.28 | 7.35±2.97 | 38.26% | 40.92% | 22.84% |
| DynamicBind (9) | 3.07±1.29 | 1.94±1.32 | 3.44±2.01 | 40.26% | 60.25% | 49.16% |
| DiffBindFR (1) | 2.38±1.00 | 0.50±0.07 | 2.11±0.80 | 63.77% | 70.40% | 59.28% |
| SurfDock (10) | 1.96±0.11 | 0.46±0.44 | 2.17±1.10 | 66.10% | 72.11% | 63.70% |
| PackDock (11) | 2.80±1.34 | 2.43±1.50 | 3.03±2.35 | 54.34% | 66.53% | 53.01% |
| FlexPose (12) | 3.01±1.65 | 2.57±0.81 | 3.59±1.56 | 41.42% | 61.64% | 31.52% |
| **CoupleDiff** | **1.62±0.43** | **0.23±0.53** | **1.82±0.74** | **69.55%** | **73.76%** | **68.63%** |

(↑) / (↓) denotes higher / lower is better; Top 1 and Top 2 are highlighted with **bold** and underlined, respectively.

and structural inaccuracies, the model demonstrates strong robustness, indicating that it can effectively integrate learned priors with imperfect structural inputs.

Overall, these results demonstrate that CoupleDiff provides a unified and robust solution for target–drug structure reconstruction, achieving consistent improvements across datasets of varying difficulty and evaluation settings. The method maintains strong performance not only on standard curated benchmarks but also under more challenging scenarios, including apo structures and predicted target backbones, highlighting its stability and generalization capability. Importantly, the observed gains are not limited to a single metric or specific component, but are consistently reflected across overall, side-chain, and ligand reconstruction quality, as well as success rate metrics. This indicates that CoupleDiff is able to produce structurally coherent and physically plausible binding conformations in a reliable manner.

**B. Results of Cross-docking Task.** We further evaluate the performance of different methods on the more challenging cross-docking task, where drugs are docked into non-cognate target structures. As shown in Table S11, CoupleDiff achieves the highest success rates on all six benchmarks, including CDK2, EGFR, and the more diverse cross-docking splits (DUDE27-HoloEns (D-H), C-D (CASF2016-Drug ), PDBbind crossdocked core set (P-CD), and ANN), demonstrating its strong ability to generalize across different target-drug combinations. At the same time, it also yields the lowest RMSD, indicating improved structural [illegible] predicted binding poses. Compared with existing methods, which exhibit performance variability across da[illegible]ets, CoupleDiff maintains consistently strong results under all evaluation scenarios. This robustness is especially notable in more challenging settings such as P-CD and ANN, where structural discrepancies between training and testing complexes are more pronounced.

**Table S7. Performance of different methods in reconstructing complex structures on the APObind set, evaluated by RMSD (Å) and success rate.**

| Method | RMSD (↓) | | | Success Rate (↑) | | |
|---|---|---|---|---|---|---|
| | Overall | Side-chain | Ligand | Overall RMSD < 2 Å | sc-RMSD < 2 Å | L-RMSD < 2 Å |
| Glide (3) | 11.55±5.11 | 9.81±4.55 | 15.68±5.21 | 17.22% | 18.50% | 9.13% |
| RosettaLigand (4) | 10.15±4.33 | 9.55±4.31 | 19.88±5.01 | 19.81% | 20.11% | 7.25% |
| AutoDock (5) | 8.95±4.91 | 7.44±4.22 | 13.81±4.77 | 20.15% | 21.33% | 11.08% |
| TankBind (6) | 5.88±3.41 | 4.95±3.51 | 15.11±5.33 | 21.43% | 27.05% | 10.19% |
| DiffDock (7) | 5.11±3.01 | 3.88±1.55 | 10.22±4.33 | 23.18% | 35.22% | 16.55% |
| DiffDock-Pocket (8) | 3.55±1.58 | <u>2.55±1.71</u> | 6.81±4.55 | 37.05% | 55.11% | 23.48% |
| DynamicBind (9) | 3.33±2.01 | 2.56±0.78 | 4.16±2.88 | <u>50.51%</u> | <u>58.22%</u> | 49.33% |
| DiffBindFR (1) | <u>3.25±1.55</u> | 2.81±0.44 | <u>3.99±1.45</u> | 48.77% | 43.18% | <u>50.04%</u> |
| SurfDock (10) | 4.88±2.51 | 2.77±1.55 | 4.99±2.69 | 46.11% | 58.01% | 44.25% |
| PackDock (11) | 3.75±1.58 | 3.71±0.45 | 4.01±1.58 | 44.33% | 46.22% | 36.11% |
| FlexPose (12) | 3.66±1.01 | 2.66±0.41 | 4.88±2.44 | 38.22% | 49.05% | 28.18% |
| **CoupleDiff** | **2.52±2.01** | **2.48±1.22** | **3.44±1.61** | **53.15%** | **60.88%** | **51.45%** |

(↑) / (↓) denotes higher / lower is better; Top 1 and Top 2 are highlighted with **bold** and <u>underlined</u>, respectively.

**Table S9. Performance of different methods in reconstructing complex structures on the ESMFold-predicted set, evaluated by RMSD (Å) and success rate.**

| Method | RMSD (↓) | | | Success Rate (↑) | | |
|---|---|---|---|---|---|---|
| | Overall | Side-chain | Ligand | Overall RMSD < 2 Å | sc-RMSD < 2 Å | L-RMSD < 2 Å |
| Glide (3) | 12.15±5.21 | 10.05±4.66 | 16.22±5.15 | 16.55% | 17.81% | 8.99% |
| RosettaLigand (4) | 10.50±4.45 | 9.88±4.41 | 20.15±5.11 | 18.88% | 19.50% | 6.95% |
| AutoDock (5) | 9.25±5.01 | 7.81±4.33 | 14.05±4.88 | 19.95% | 21.10% | 11.24% |
| TankBind (6) | 6.11±3.55 | 5.02±3.61 | 15.33±5.40 | 21.78% | 27.55% | 9.88% |
| DiffDock (7) | 4.22±3.15 | 4.05±1.68 | 10.51±4.44 | 33.50% | 45.18% | 16.21% |
| DiffDock-Pocket (8) | 3.68±1.77 | 4.65±1.88 | 7.01±4.55 | 36.95% | 44.88% | 23.99% |
| DynamicBind (9) | <u>3.45±2.11</u> | 4.71±0.85 | 3.88±2.91 | 36.21% | 28.55% | 39.02% |
| DiffBindFR (1) | 4.25±1.66 | 6.88±0.51 | 4.05±1.55 | 28.81% | 22.95% | 32.95% |
| SurfDock (10) | 3.99±2.65 | 3.79±1.61 | 4.15±2.81 | <u>46.80%</u> | <u>48.15%</u> | <u>44.05%</u> |
| PackDock (11) | 3.95±1.69 | <u>3.75±0.55</u> | <u>3.65±1.66</u> | 39.11% | 36.05% | 43.18% |
| FlexPose (12) | 4.88±3.11 | 5.65±0.48 | 3.95±2.45 | 30.20% | 28.99% | 38.15% |
| **CoupleDiff** | **3.45±2.05** | **3.21±1.25** | **3.58±1.68** | **48.15%** | **49.20%** | **45.40%** |

(↑) / (↓) denotes higher / lower is better; Top 1 and Top 2 are highlighted with **bold** and <u>underlined</u>, respectively.

**Table S11. Performance of different methods on various cross-docking datasets, evaluated by overall success rate and overall RMSD (Å).**

| Method | Overall Success Rate (%) ↑ | | | | | | Overall RMSD (Å) ↓ | | | | | |
|---|---|---|---|---|---|---|---|---|---|---|---|---|
| | CDK2 | EGFR | D-H | C-D | P-CD | ANN | CDK2 | EGFR | D-H | C-D | P-CD | ANN |
| AutoDock (5) | 5.92 | 3.11 | 11.37 | 18.47 | 27.88 | 15.29 | 6.12 | 6.37 | 6.04 | 4.37 | 4.89 | 55.37 |
| Glide (3) | 15.17 | 9.38 | 13.10 | 16.32 | 40.64 | 13.38 | 5.21 | 6.87 | 5.72 | 4.94 | 4.12 | 4.86 |
| DiffDock (7) | 29.83 | 33.55 | <u>34.78</u> | 32.72 | 43.92 | 21.40 | 4.06 | 5.46 | 3.17 | 3.46 | 2.85 | 4.02 |
| TankBind (6) | 45.20 | <u>41.83</u> | 31.69 | 20.98 | 41.37 | 24.01 | 2.17 | <u>2.21</u> | <u>3.14</u> | 4.10 | 3.05 | 3.90 |
| DiffBindFR (1) | 39.45 | 41.05 | 34.24 | 63.02 | 56.24 | <u>54.56</u> | 1.85 | 2.58 | 3.28 | 2.23 | 1.80 | <u>2.43</u> |
| DiffDock-Pocket (8) | 44.28 | 30.75 | 29.05 | 40.81 | 50.37 | 30.12 | 1.82 | 5.50 | 3.91 | 3.12 | 1.93 | 3.11 |
| PackDock (11) | <u>50.64</u> | 28.37 | 20.46 | 61.13 | 61.45 | 42.63 | <u>1.73</u> | 4.02 | 4.56 | 2.31 | 1.84 | 2.94 |
| FlexPose (12) | 48.56 | 33.22 | 26.81 | <u>69.24</u> | <u>69.33</u> | 36.74 | 1.80 | 3.13 | 5.13 | <u>1.38</u> | <u>1.66</u> | 3.05 |
| **CoupleDiff** | **56.01** | **43.46** | **40.52** | **70.06** | **76.58** | **60.85** | **1.72** | **2.14** | **2.80** | **1.05** | **1.57** | **2.28** |

(↑) / (↓) denotes higher / lower is better; Top 1 and Top 2 are highlighted with **bold** and <u>underlined</u>, respectively.

**C. Ablation Study.** To evaluate the contributions of key components in our model, we conducted ablation experiments by systematically removing pre-docking strategy, physical features, cooperative scalar-vector representation module, target embedding discretization module, physics-constrained interaction module, diffusion module and physical constraints, as illustrated in Table S13.

**Criticality of the Pre-Docking Strategy.** The pre-docking strategy initializes drug poses to avoid local energy minima. Adding a pre-docking strategy improves the drug success rate by 3.14%, demonstrating that accurate geometric initialization acts synergistically with structural modeling to preserve global conformational fidelity, preventing pose divergence during refinement.

**Importance of Physical Features.** Physical features (e.g., atomic mass, charge) offer intrinsic molecular descriptors. Removing them caused a notable rise in sc-RMSD from 0.29Å to 0.31Å and a decrease in drug success rate by 1.04%, which highlights that explicit physical priors enhance local structural alignment and enhance atomic-level discriminability.

**Effectiveness of the Cooperative Scalar-Vector Representation Module.** The cooperative scalar-vector representation module is designed to model the structural interdependencies. Removing the cooperative scalar-vector representation module significantly degraded side-chain prediction accuracy, increasing RMSD from 0.29Å to 2.04Å, while the drug success rate dropped by 17.47%, highlighting the critical importance of this module in maintaining accurate structural predictions.

**Role of the Target Embedding Discretization Module.** Discretization encodes local geometric details at fine resolution. Its removal resulted in a substantial degradation: the overall RMSD increased from 2.01Å to 3.34Å, and the success rate dropped from 61.60% to 41.34%. These results emphasize that discretized spatial encoding is indispensable for capturing spatial granularity critical for precise structure generation.

**Effect of the Physics-Constrained Interaction Module.** Physics-constrained interactions impose inter-molecular force priors. Eliminating this component increased the overall RMSD from 2.01Å to 2.11Å and reduced drug success rate by 8.69%. This suggests that energy-aware modeling facilitates spatial compatibility between target and drug, and valid predictions.

**Contribution of the Diffusion Module.** Diffusion modeling introduces denoising dynamics to capture conformation flexibility. Removing it led to severe performance loss, with overall RMSD increasing by 2.98Å and drug success rate dropping by 30.79%. This underlines that the generative denoising process is fundamental for modeling complex structural distributions in a physically consistent manner.

**Necessity of Physical Constraints.** Physical constraints include seven physical and geometrical terms to ensure structural plausibility. Adding these constraints significantly mitigates deviations, the overall RMSD decreases by 0.79Å, and overall success rate rises by 14.76%, which demonstrates that physical constraints prioritize local atomic realism over global sampling, leveraging physicochemical principles to refine modeling.

**Table S13. Experimental results on ablation study, which summarizes the performance of models with various components (✓/✗), highlighting the impact of each element. a: pre-docking strategy, b: physical features, c: cooperative scalar-vector representation module, d: target embedding discretization module, e: physics-based interaction module, f: diffusion module, g: physical constraints.**

| Components | | | | | | | RMSD ↓ | | | Success Rate ↑ | | |
|---|---|---|---|---|---|---|---|---|---|---|---|---|
| a | b | c | d | e | f | g | Overall | Side-chain | Ligand | Overall RMSD <2Å | sc-RMSD <2Å | L-RMSD <2Å |
| ✓ | ✓ | ✓ | ✓ | ✓ | ✓ | ✓ | **2.01 Å** | **0.29 Å** | **2.17 Å** | **65.30%** | **70.50%** | **61.60%** |
| ✗ | ✓ | ✓ | ✓ | ✓ | ✓ | ✓ | 2.07 Å | 0.48 Å | 2.31 Å | 64.17% | 69.17% | 58.46% |
| ✓ | ✗ | ✓ | ✓ | ✓ | ✓ | ✓ | 2.05 Å | 0.31 Å | 2.73 Å | 64.66% | 70.15% | 60.56% |
| ✓ | ✓ | ✗ | ✓ | ✓ | ✓ | ✓ | 3.22 Å | 2.04 Å | 4.96 Å | 49.72% | 62.01% | 44.13% |
| ✓ | ✓ | ✓ | ✗ | ✓ | ✓ | ✓ | 3.34 Å | 1.89 Å | 5.13 Å | 47.01% | 67.88% | 41.34% |
| ✓ | ✓ | ✓ | ✓ | ✗ | ✓ | ✓ | 2.11 Å | 0.59 Å | 2.70 Å | 60.87% | 57.45% | 52.91% |
| ✓ | ✓ | ✓ | ✓ | ✓ | ✗ | ✓ | 4.99 Å | 3.72 Å | 4.08 Å | 40.45% | 36.67% | 30.81% |
| ✓ | ✓ | ✓ | ✓ | ✓ | ✓ | ✗ | 2.80 Å | 1.93 Å | 5.42 Å | 50.54% | 66.21% | 43.02% |
| ✗ | ✓ | ✗ | ✓ | ✓ | ✓ | ✓ | 4.53 Å | 3.65 Å | 3.81 Å | 41.08% | 38.90% | 35.27% |
| ✓ | ✓ | ✗ | ✓ | ✓ | ✓ | ✗ | 4.60 Å | 3.06 Å | 4.26 Å | 37.19% | 37.12% | 30.78% |
| ✗ | ✓ | ✗ | ✓ | ✓ | ✓ | ✗ | 5.01 Å | 5.17 Å | 6.11 Å | 29.23% | 25.45% | 26.09% |

(↑) / (↓) denotes higher / lower is better; Top 1 and Top 2 are highlighted with **bold** and underlined, respectively.

**D. Quantitative Benchmarking in Large-Scale Virtual Screening and Binding Affinity Prediction.** To demonstrate the robustness and predictive power of the representations learned by CoupleDiff, we evaluated its performance on two of the most important tasks in SBDD: large-scale virtual screening and binding affinity prediction.

To evaluate the practical utility in hit identification, we performed large-scale virtual screening on the widely-used DUD-E and DEKOIS2.0 benchmarks. A model's performance in this task is defined by two distinct capabilities: its overall ranking accuracy across the entire dataset, measured by area under the receiver operating characteristic curve (AUC), and its efficiency in early hit retrieval, measured by relative enrichment (RE).

**Table S15. Performance of different methods in virtual screening on DUD-E dataset, evaluated by AUC and RE at various percentages.**

| Method | AUC ↑ | RE 0.5% ↑ | RE 1.0% ↑ | RE 2.0% ↑ | RE 5.0% ↑ |
|---|---|---|---|---|---|
| Glide (3) | 0.602 | 5.491% | 3.629% | 2.102% | 1.264% |
| RosettaLigand (4) | 0.658 | 10.351% | 8.451% | 5.037% | 2.394% |
| AutoDock(5) | 0.620 | 9.139% | 7.290% | 4.341% | 2.073% |
| TankBind(6) | 0.713 | 11.462% | 6.025% | 3.026% | 2.563% |
| DiffDock(7) | 0.744 | 43.275% | 26.487% | 19.341% | 10.375% |
| DiffDock-Pocket(8) | 0.806 | 50.861% | 34.243% | 22.241% | 12.072% |
| DynamicBind(9) | 0.935 | 83.529% | 45.874% | 30.952% | 14.561% |
| DiffBindFR(1) | 0.951 | <u>101.462%</u> | 60.926% | 35.318% | 13.451% |
| SurfDock(10) | 0.947 | 96.351% | 50.648% | 30.592% | 12.922% |
| PackDock(11) | 0.960 | 99.460% | 58.492% | 33.296% | 13.647% |
| FlexPose(12) | <u>0.966</u> | 100.518% | <u>61.395%</u> | <u>35.793%</u> | <u>15.302%</u> |
| **CoupleDiff** | **0.972** | **107.471%** | **63.284%** | **38.351%** | **17.462%** |

(↑) / (↓) denotes higher / lower is better; Top 1 and Top 2 are highlighted with **bold** and <u>underlined</u>, respectively.

**Table S16. Performance of different methods in virtual screening on DEKOIS2.0 dataset, evaluated by AUC and RE at various percentages.**

| Method | AUC ↑ | RE 0.5% ↑ | RE 1.0% ↑ | RE 2.0% ↑ | RE 5.0% ↑ |
|---|---|---|---|---|---|
| Glide (3) | 0.512 | 4.127% | 2.819% | 1.543% | 0.966% |
| RosettaLigand (4) | 0.541 | 7.831% | 6.105% | 3.829% | 1.811% |
| AutoDock(5) | 0.525 | 6.518% | 5.244% | 3.167% | 1.582% |
| TankBind(6) | 0.603 | 9.176% | 4.591% | 2.238% | 1.903% |
| DiffDock(7) | 0.624 | 33.722% | 19.851% | 14.570% | 7.845% |
| DiffDock-Pocket(8) | 0.686 | 40.109% | 25.663% | 16.792% | 9.127% |
| DynamicBind(9) | 0.785 | 65.234% | 34.488% | 23.215% | 11.059% |
| DiffBindFR(1) | 0.811 | 79.547% | 45.716% | 26.539% | 10.198% |
| SurfDock(10) | 0.802 | 75.882% | 38.023% | 22.951% | 9.776% |
| PackDock(11) | <u>0.820</u> | 80.165% | 44.978% | 25.084% | 10.332% |
| FlexPose(12) | 0.815 | <u>81.438%</u> | <u>46.125%</u> | <u>27.804%</u> | <u>11.521%</u> |
| **CoupleDiff** | **0.832** | **85.642%** | **48.219%** | **29.133%** | **13.107%** |

(↑) denotes higher lower is better; Top 1 and Top 2 are highlighted with **bold** and <u>underlined</u>, respectively.

On the DUD-E benchmark (Supplementary Tab. C7), CoupleDiff established its SOTA performance with a top-ranked AUC of 0.972, confirming its globally accurate ranking ability, as shown in the distribution in Fig. S1a. More importantly, it demonstrated exceptional early enrichment. The RE of 107.5% at the stringent 0.5% threshold is a crucial finding in Fig. S1b, as this signifies an over 100-fold enrichment of true binders compared to random selection. The efficiency in prioritizing active compounds at the top of the ranked list directly translates into reduced experimental cost and time. Such performance stems from the model's adaptive design, which enables CoupleDiff to capture subtle, high-affinity interactions characteristic of true binders, thereby consistently ranking them among the top candidates.

To validate findings on a more diverse and challenging chemical space, we then evaluated all methods on the DEKOIS2.0 benchmark (Supplementary Tab. C8). While this dataset proved more difficult for all models, CoupleDiff maintained its decisive advantage. It again secured the highest AUC (0.832) in Fig. S1c and consistently led across all RE metrics in Fig. S1d, including a top RE of 85.6% at 0.5%. The stable performance across both benchmarks suggests that the learned representation captures transferable biophysical principles rather than dataset-specific heuristics, demonstrating robustness to variations in chemical space and target composition.

Taken together, the top-tier performance on both benchmarks demonstrates a critical dual capability: CoupleDiff not only possesses a globally accurate understanding of binding affinity but also excels at the vital and practical task of efficient early hit identification. This combination of global accuracy and early enrichment efficiency establishes CoupleDiff as a robust and highly effective tool for real-world virtual screening campaigns.

We then evaluated CoupleDiff on the downstream binding affinity prediction task, comparing it against a diverse set of structure-based deep learning models. As shown in Fig. S1e-l and summarized in Supplementary Tab. S17, CoupleDiff achieved the strongest overall performance, with the highest Pearson correlation coefficient (PCC = 0.820) and the lowest root mean square error (RMSE = 1.20). This superior performance indicates that the model has learned a quantitatively accurate mapping between structure and binding strength, rather than merely separating binders from non-binders. We attribute this advantage to CoupleDiff's joint modeling of ligand pose optimization and receptor adaptation, which enables the network to capture subtle, drug-specific energetic contributions arising from induced fit and local structural rearrangements within the binding pocket.

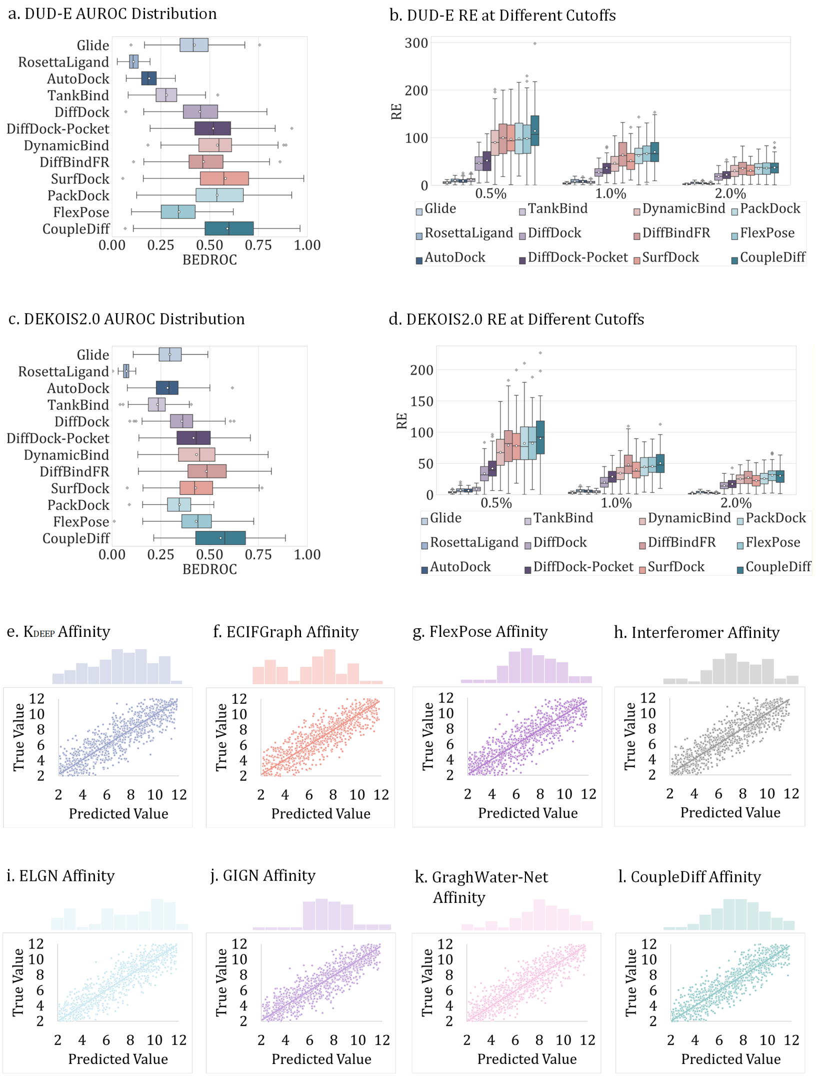


**Fig. S1. Comparative performance analysis in large-scale virtual screening benchmarks and binding affinity quantification. a, b** AUC distribution and early hit identification efficiency at different selection thresholds (0.5%, 1.0%, and 2.0%) of all evaluated models on DUD-E dataset. **c, d** AUC distribution reflecting the global ranking precision of all evaluated models and early hit identification efficiency at different selection thresholds (0.5%, 1.0%, and 2.0%) of all evaluated models on DEKOIS2.0 dataset. **e-l** Binding affinity correlation comparing predicted values and experimental ground truths for multiple methods.

Across both virtual screening and affinity prediction tasks, CoupleDiff consistently demonstrates strong generalization and quantitative accuracy. The ability to identify true binders at early screening stages while providing reliable affinity estimates provides compelling evidence that the learned representation encodes meaningful biophysical information underlying molecular recognition, rather than relying on superficial statistical correlations.

**E. Imatinib Selectivity Across Homologous Tyrosine Kinase Family.** To assess whether CoupleDiff can discriminate drug selectivity under highly challenging structural conditions, we conducted a detailed case study on Iimatinib, a prototypical type-II tyrosine kinase inhibitor with a well-established but highly selective target ensemble. imatinib potently inhibits ABL1 and c-Kit kinases, while exhibiting little to no activity against closely related kinases such as EGFR and VEGFR2 (19). Importantly, these kinases share a conserved fold and highly similar inactive-state conformations, making selectivity prediction based solely on receptor structure intrinsically difficult.

We selected several homologous protein kinases representing the full spectrum of imatinib's activity: the primary on-target, human ABL1 kinase (UniProt: P00519), using three distinct wild-type structures (PDB IDs: 1IEP, 2HYY, 3GVU) as positive controls; a key off-target, human c-Kit kinase (UniProt: P10721, PDB: 1T46), as a secondary positive control; and two non-targets, human EGFR (UniProt: P00533, PDB: 2GS6) and human VEGFR2 (UniProt: P35968, PDB: 4ASD), as crucial negative controls. As shown in Fig. S2b, pairwise RMSD analysis of the apo binding pockets reveals a high degree of structural similarity across all six kinases prior to drug binding. This pronounced structural homogeneity highlights the intrinsic difficulty of the task: prior to interaction with imatinib, target and non-target kinases are nearly indistinguishable based on geometry alone.

**Table S17. Performance of different methods in predicting target-drug binding affinity, evaluated by PCC, RMSE, Spearman, and MAE.**

| Method | PCC ↑ | RMSE ↓ | Spearman ↑ | MAE ↓ |
|---|---|---|---|---|
| $K_{DEEP}$ (13) | 0.734 | 1.50 | 0.725 | 1.12 |
| ECIFGraph::HM-Holo-Apo (14) | 0.781 | 1.42 | 0.765 | 1.08 |
| FlexPose (12) | 0.793 | 1.39 | 0.780 | 1.06 |
| Interformer (15) | 0.802 | <u>1.27</u> | 0.790 | 1.04 |
| ELGN (16) | 0.805 | 1.31 | 0.795 | 1.07 |
| GIGN (17) | 0.807 | 1.34 | 0.800 | 1.10 |
| GraphWater-Net (18) | <u>0.814</u> | 1.28 | <u>0.810</u> | <u>1.05</u> |
| **CoupleDiff** | **0.820** | **1.20** | **0.815** | **1.00** |

(↑) / (↓) denotes higher / lower is better; Top 1 and Top 2 are highlighted with **bold** and <u>underlined</u>, respectively.

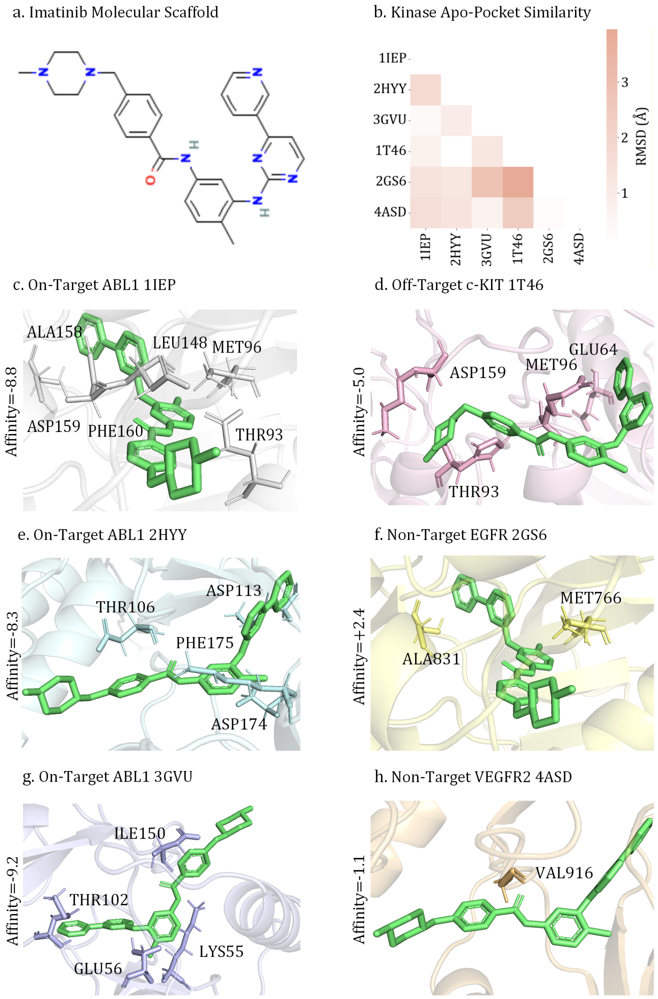


**Fig. S2. Structural determinants of drug selectivity in the homologous tyrosine kinase family. a** Imatinib molecular scaffold: The chemical identity of imatinib. **b** Kinase apo-pocket similarity: Pairwise RMSD analysis showing high structural homogeneity across all six kinase binding pockets prior to drug binding. **c, e, g** Predicted binding poses of on-target ABL1 targets (1IEP, 2HYY, 3GVU) binding of imatinib. **d** Off-target c-Kit 1T46: Predicted binding pose of 1T46 and imatinib. **f** Non-target EGFR 2GS6: Predicted binding pose of 2GS6 and imatinib. **h** Non-target VEGFR2 4ASD: Predicted binding pose of 4ASD and imatinib.

Despite the high degree of pre-binding structural similarity, CoupleDiff successfully differentiates targets from non-targets once the drug-target interaction is explicitly modeled. As illustrated in Fig. S2c-h, imatinib forms stable, low-energy complexes with all three ABL1 structures, with predicted affinities ranging from −8.3 to −9.2 kcal/mol, and maintains favorable binding to c-Kit (−5.0 kcal/mol). In contrast, substantially weakened or unfavorable binding is predicted for EGFR (+2.4 kcal/mol) and VEGFR2 (−1.1 kcal/mol), correctly identifying them as non-targets despite their structural resemblance to ABL1.

Having established its functional fidelity, we next assessed the structural accuracy of the predicted binding poses. Structural inspection of the predicted complexes reveals that this discrimination arises from target-specific incompatibilities that only emerge upon drug engagement. In ABL1 and c-Kit, imatinib adopts the canonical type-II binding mode, extending into the hydrophobic back pocket and forming a conserved network of stabilizing interactions in the inactive kinase conformation (Fig. S2c,d,e,g). These binding geometries resemble experimentally observed complexes, indicating that the pocket architecture of these kinases is intrinsically compatible with imatinib's scaffold.

In contrast, distinct structural failure modes are observed for the non-target kinases. For the EGFR target, the predicted binding mode reveals the unfavorable interactions originating from a primary steric conflict (Figure S2f). The core scaffold of imatinib clashes severely with the side chain of MET766. This bulky methionine replaces the much smaller valine found at the equivalent position in ABL1, physically obstructing the binding pocket and preventing the inhibitor from achieving its optimal orientation. This initial misplacement triggers a secondary and equally critical failure: the altered orientation of imatinib prevents its amide group from forming the key hydrogen bond with the backbone of the catalytic loop residue, Aspartate 831 (ALA831). The loss of this interaction, which is essential for stabilizing the DFG-out conformation in ABL1, further destabilizes the entire complex.

For the VEGFR2 target, the model identified a distinct but equally prohibitive structural barrier (Figure S2h). Here, the primary issue is the larger gatekeeper residue, VAL916, which is a threonine in ABL1. This hydrophobic residue constricts the entrance to the deep hydrophobic pocket essential for imatinib's characteristic DFG-out binding configuration. The model predicts that this steric obstruction not only prevents the inhibitor from fully engaging with the pocket but also abrogates the crucial hydrogen bond that imatinib forms with the gatekeeper's backbone in ABL1.

Together, these results demonstrate that CoupleDiff can resolve drug selectivity not by relying on pre-existing structural differences among homologous targets, but by explicitly modeling how a drug perturbs and interrogates each binding pocket. Even when targets are highly similar prior to binding, the model correctly identifies subtle incompatibilities that govern selective inhibition, such as a steric and electrostatic failure in EGFR or a gatekeeper-blockade in VEGFR2. This capacity to infer selectivity from drug-induced structure is essential for SBDD within closely related protein families.

**F. Identification and Validation of Novel Hit Scaffolds for HIV-1 Protease.** Having established the performance of CoupleDiff across a range of benchmarks, we next validated its practical utility in more closely mimic real-world drug discovery challenges. We selected HIV-1 protease as a representative target, which plays a pivotal role in the HIV viral replication cycle (20), as its inhibition directly blocks the formation of infectious viral particles by disrupting viral maturation, while decades of research have yielded numerous clinically approved inhibitors and over 5,000 high-resolution crystal structures, including apo states, ligand-bound complexes, and mutant variants, that provide a robust experimental foundation for simulation validation (21). Additionally, its unique structural dynamics, as a homodimer with two flexible flap regions flanking the active site, undergo significant conformational changes from an open to a closed state upon ligand binding, an induced-fit effect that MD simulations excel at capturing whereas traditional static docking struggles to accurately characterize (22). Furthermore, HIV-1 readily develops drug resistance through protease mutations (e.g., D30N, V82A) that alter the binding pocket's shape and electrostatics, presenting a stringent test for evaluating the applicability of simulation tools in complex clinical scenarios (23). Coupled with its modest molecular weight ($\sim$22 Da), it achieves a balance between computational feasibility, which allows long-duration all-atom simulations, and strong validation potential (24). The flexible and polar active site, together with the challenges introduced by drug-resistant mutations, provides a comprehensive assessment of whether screening and docking pipelines can handle dynamic targets and identify novel and effective chemical scaffolds.

***F.1. Virtual Screening and Candidate Selection.*** The overall workflow of hit identification is illustrated in Fig. S3a. The screening process was initiated from the ZINC15 database, a library containing over six million drug-like protomers enriched with compounds in the desirable molecular weight range of 250-450 Da and a LogP between one and four, making it an ideal starting point for hit identification (25). The first step was a large-scale 2D similarity search, using four clinically approved HIV-1 protease inhibitors as query structures including lopinavir, darunavir, saquinavir, and ritonavir. These query molecules represent diverse chemical scaffolds and well-characterized binding modes to the target protein, ensuring broad coverage of biologically relevant structural space. To quantify molecular similarity, Morgan fingerprints were employed for structural characterization. For every valid candidate molecule, each one was structurally parsed, with invalid structures excluded to ensure data quality. For each valid candidate, a corresponding fingerprint was generated, enabling consistent structural comparison. The Tanimoto similarity coefficient, a standard metric for evaluating molecular structural overlap, ranging from 0 (no similarity) to 1 (identical structures), was calculated between each candidate's fingerprint and those of the four query inhibitors. The highest similarity score across all queries was retained to assess its maximum structural relatedness to known inhibitors. A strict Tanimoto similarity threshold of 0.3 was adopted to filter for novel scaffolds: only candidates with a highest similarity score $\leq$ 0.3 were retained as hits. This threshold was selected to effectively exclude compounds sharing significant structural overlap with known inhibitors, while prioritizing molecules with unique substructural features that diverge from clinically validated scaffolds. This strategy minimizes the risk of re-discovering molecules with similar binding modes to existing inhibitors and maximizes the potential for identifying novel chemical entities that may interact with HIV-1 protease. Through this targeted screening, the original library of six million molecules was efficiently refined into a set of 1,040 promising candidates, which directly facilitated the subsequent docking step, streamlining the workflow by prioritizing molecules with a high potential for structural novelty. These candidates were then docked to the HIV-1 protease target using CoupleDiff to generate 1,040 protein-ligand complex structures. Based on the predicted binding scores, the top 24 molecules were selected for validation via all-atom MD simulations.

From these candidates, six compounds were prioritized as final hits. Their 3D chemical structures are shown in Fig. S3b-g, highlighting their chemical diversity and structural distinctness from existing HIV-1 protease inhibitors. Despite their low 2D similarity to known drugs, all six compounds were predicted to bind within the canonical substrate-binding pocket.

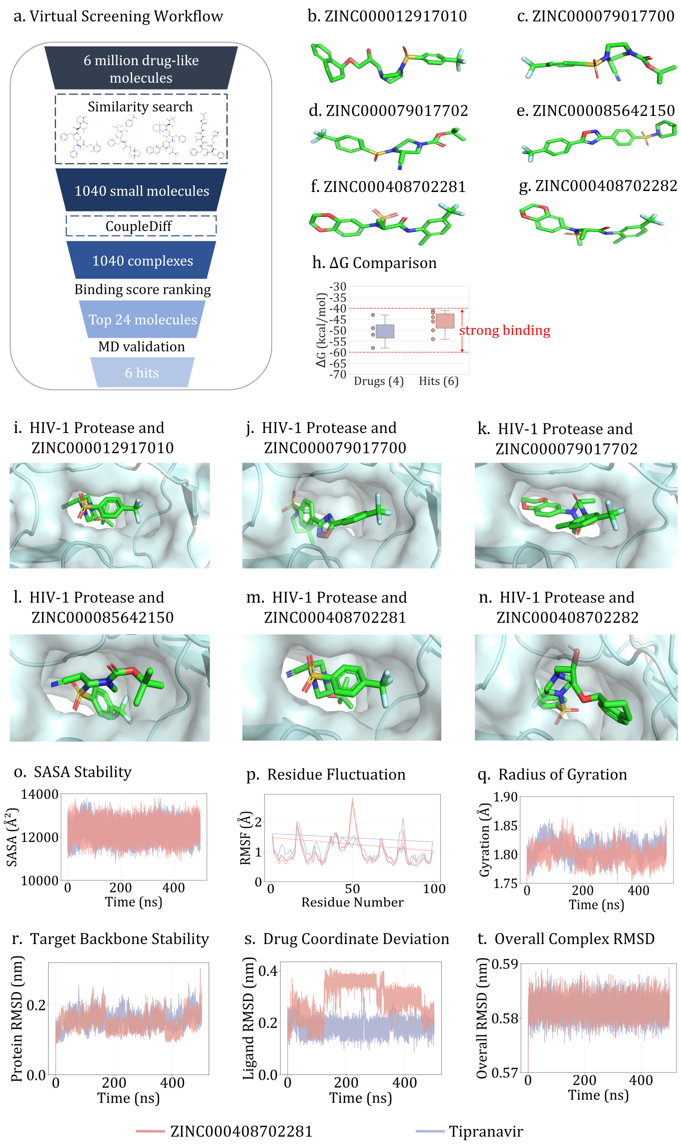


**Fig. S3. Identification and validation of novel therapeutic scaffolds for HIV-1 Protease. a** Virtual screening workflow: Screening pipeline starting from six million molecules to prioritize potential inhibitors through similarity search and docking. **b-g** Structural diversity of six hit scaffolds highlighting distinct chemical identities and low similarity to known drugs. **h** ΔG comparison: Binding free energy between the identified hits and four clinically approved inhibitors. **i-n** Predicted binding poses in the active site depicting stabilized drug orientations and specific pocket engagement. **o-t** Dynamic stability analysis from a 500ns MD simulation comparing the hit ZINC000408702281 with the reference drug tipranavir.

***F.2. Energetic and Structural Validation.*** To further assess the thermodynamic favorability of these interactions, binding free energies ($\Delta G$) were estimated and compared with those of four clinically approved HIV-1 protease inhibitors (Fig. S3h). Consistent with the stable docking geometries, several hit compounds exhibited binding free energies comparable to the reference drugs, indicating that scaffold novelty was not achieved at the expense of thermodynamic stability. These results suggest that CoupleDiff can prioritize chemically distinct compounds that form energetically favorable complexes with the target protein.

The corresponding docking poses of the six hit compounds are shown in Fig. S3i-n. Each compound adopts a well-defined binding orientation within the active site, forming favorable interactions with key catalytic residues and the flexible flap regions. The diversity of binding modes observed across the six hits indicates that CoupleDiff can predict multiple viable interaction patterns while preserving essential target engagement, underscoring its ability to identify structurally novel scaffolds beyond established chemotypes.

***F.3. Mechanistic Validation via MD Simulations.*** The dynamic stability of the predicted complexes was then evaluated using 500ns all-atom MD simulations. A representative top hit, ZINC000408702281, was selected for direct comparison with the clinically approved inhibitor tipranavir. Multiple structural metrics were monitored throughout the simulations to characterize global and local stability (Fig. S3o-t). The solvent-accessible surface area (SASA) and radius of gyration remained stable over time (Fig. S3o, q), indicating preserved global compactness and solvation behavior. Per-residue RMSF analysis revealed limited fluctuations in the binding pocket residues, comparable to those observed for tipranavir (Fig. S3p). The protein backbone RMSD rapidly converged and remained below 0.30 nm throughout the simulation (Fig. S3r), confirming global structural stability.

Importantly, ligand RMSD values, calculated after alignment to the protein backbone, remained tightly constrained within a narrow range (<0.25 nm) (Fig. S3s), while the overall complex RMSD further confirmed long-term stability of the protein-ligand assembly (Fig. S3t).

Collectively, by identifying chemically novel scaffolds that form stable and energetically favorable complexes with a highly dynamic and clinically relevant target, CoupleDiff bridges the gap between high-throughput virtual screening and rigorous mechanistic validation. This capability highlights its potential as a practical tool for discovering novel hits against challenging drug targets in real-world drug discovery.

**G. Adaptive Weight Adjustment.** In MTL frameworks, the design of loss weightings critically influences how the model balances its optimization trajectory across competing objectives. In our model, the total training loss comprises three core components: the structural loss $\mathcal{L}_{\text{structure}}$, the diffusion denoising loss $\mathcal{L}_{\text{noise}}$, and the physics and geometry-guided loss $\mathcal{L}_{\text{phy-geo}}$, jointly supervising the complex modeling.

Traditionally, scalar weights are statically assigned to each loss component. However, such static weighting cannot adapt to the non-stationary nature of training. Different tasks may converge at different rates or present varying levels of difficulty throughout training. Improper weighting may result in biased gradient updates, overfitting to easier tasks, or neglecting harder ones. To overcome this, we introduce a dynamically adaptive inverse proportional weighting strategy based on instantaneous loss magnitudes.

***G.1. Theoretical Motivation.*** From a theoretical point of view, our strategy can be interpreted under a Min-Max loss balancing framework. Specifically, we seek to minimize the largest normalized task loss at each training step, formulated as:

$$\max_{i\in\{s,d,p\}} \left(\frac{\ell_i}{\alpha_i}\right) \tag{22}$$

where $\ell_i$ denotes the instantaneous scalar loss,$\alpha_i$ represents the corresponding adaptive weights.

Under our inverse proportional weighting rule, this ratio is approximately constant across all loss terms:

$$\frac{\ell_i}{\alpha_i} \approx \sum_j \ell_i \cdot \left(\frac{1/\ell_j}{\sum_k 1/\ell_k}\right) = \text{const.} \tag{23}$$

This normalization implies that the optimizer perceives all tasks as having equal scaled difficulty, thus promoting fairness in multi-objective convergence and avoiding task imbalance.

***G.2. Formulation of Inverse Proportional Normalization.*** At each training step $t$, we observe the instantaneous (i.e., per-batch) losses:

$$\mathcal{L}^{(t)}_{\text{structure}} = \ell_s, \quad \mathcal{L}^{(t)}_{\text{diffusion}} = \ell_d, \quad \mathcal{L}^{(t)}_{\text{phy-geo}} = \ell_p. \tag{24}$$

where $\ell_s, \ell_d, \ell_p \in \mathbb{R}_{>0}$ are the current magnitudes of the structure, diffusion, and physical geometry losses, respectively.

We then define the adaptive weights using the inverse proportional rule:

$$\alpha_i = \frac{1/\ell_i}{\sum_{j\in\{s,d,p\}} 1/\ell_j} \quad \text{for each } i \in \{s,d,p\} \tag{25}$$

This ensures that the sum of weights satisfies $\sum_i \alpha_i = 1$, while favoring tasks with smaller loss magnitudes by reducing their weight, and increasing the weight on larger-loss terms that need more attention.

The final total loss is as follows:

$$\mathcal{L}^{(t)}_{\text{total}} = \alpha_s \cdot \ell_s + \alpha_d \cdot \ell_d + \alpha_p \cdot \ell_p \tag{26}$$

where each term is adaptively scaled to counteract imbalance across sub-tasks.

***G.3. Empirical Results.*** We evaluate the impact of adaptive weighting in Table S19, comparing fixed scalar weights to our adaptive strategy. Notably, the adaptive scheme achieves the best performance across all evaluation metrics, demonstrating superior capability in fine-grained structure modeling.

**Table S19. Experimental results on different weight adjustment strategies of the loss function. a: the weight of the structure term. b: the weight of the diffusion term. c: the weight of the physical and geometry term.**

| Weight | | | RMSD ↓ | | Success Rate ↑ | |
|---|---|---|---|---|---|---|
| a | b | c | Overall | Side-chain | Drug <2Å | Overall <2Å |
| 1.0 | 1.0 | 1.0 | 2.22Å | 1.04Å | 49.13% | 51.32% |
| 0.5 | 1.0 | 1.5 | 2.43Å | 1.65Å | 45.27% | 49.48% |
| dynamic | dynamic | dynamic | **2.01Å** | **0.29Å** | **61.60%** | **65.30%** |

(↑) / (↓) denote higher / lower is better. Top 1 results are **bold**.

**H. Conformational Ensemble Docking Performance.** The heatmap in Figure S4 further emphasizes the consistency of CoupleDiff across heterogeneous targets. The L-RMSD distributions across the DUDE27 benchmark illuminate core mechanistic divergences in how docking methods address receptor flexibility, conformational selection, and induced-fit coupling. CoupleDiff achieves the lowest L-RMSD in 18 out of 27 targets. By leveraging physics-constrained optimization, CoupleDiff resolves steric conflicts and captures long-range coupling, which is critical for targets with substantial loop or helix remodeling, such as BRAF (2.70 Å), where DFG-out to DFG-in transitions are necessary for accurate binding pose prediction.

In contrast, rigid-receptor docking methods like AutoDock and Glide systematically underperform on conformationally adaptive proteins. For example, in EGFR, the kinase C-helix undergoes a pronounced shift upon ligand engagement, a transition not accounted for by static scoring protocols, resulting in a marked deviation (AutoDock: 6.36 Å vs. CoupleDiff: 3.02 Å). These methods are intrinsically limited by their reliance on a single, often non-native, receptor conformation. TankBind demonstrates reasonable performance when structural homologs are available but degrade significantly on divergent folds or allosteric pockets (e.g., GRIA2: 10.33 Å), highlighting their dependency on prior alignment and conformational priors. While DiffDock demonstrates strong performance when docking to protein conformations that closely resemble those encountered during training, its ability to generalize deteriorates in extrapolative scenarios involving rare or unseen structural variations. This limitation is exemplified by the MET target, where DiffDock achieves a relatively high L-RMSD of 7.91 Å. The underlying issue stems from the model's reliance on implicit geometric representations learned from training data, which tend to favor commonly observed backbone and side-chain configurations. As a result, DiffDock often fails to accurately model rare but functionally important conformational states, such as the loop opening events observed in MET, due to insufficient exposure to such structural diversity during training. Hybrid approaches such as DiffBindFR, which integrate learned priors with restrained physics-based refinement, show moderate success in flexible regions (e.g., CDK2: 2.71 Å) but are still limited in sites requiring long-range conformational propagation, such as TGFR1 (3.61 Å), where allosteric activation involves distal loop dynamics and solvation effects. Fragment-based flexible methods like FlexPose capture local side-chain and loop adjustments (THRB: 3.49 Å), but are insufficient for large-scale domain or backbone reorganization.

**I. Case Study on Reconstructing Accuracy.** The study of Aceclofenac binding to the 6Y3C target structure carries real-world implications for anti-inflammatory drug design. Aceclofenac is a widely used nonsteroidal anti-inflammatory drug (26), and 6Y3C corresponds to the crystal structure of cyclooxygenase-2 (COX-2), a crucial enzyme involved in prostaglandin synthesis and inflammation. Given the clinical relevance of COX-2 selectivity in minimizing gastrointestinal side effects while preserving therapeutic efficacy, accurate modeling of the binding interaction between Aceclofenac and 6Y3C is of paramount pharmaceutical interest. What makes this case particularly meaningful is due to the intricate nature of their binding mechanism, which presents significant challenges for structure reconstruction and pose prediction. Aceclofenac binds to COX-2 through a combination of hydrogen bonding, $\pi$–$\pi$ stacking, and hydrophobic interactions within a highly adaptive and conformationally flexible active site. The induced fit nature of COX-2, characterized by local rearrangements of the side chains (particularly in the hydrophobic channel and the T380, L384 and V423 residues), often leads to substantial deviations between the apo and holo states. These conformational changes, coupled with the drug's non-rigid scaffold and multiple rotatable bonds, introduce a high degree of uncertainty into conventional docking methods. By using this system, we aim to highlight our model's ability to capture such nuanced drug-induced conformational adaptations and accurately reconstruct the binding pose in a physically plausible manner, thus demonstrating its capacity to handle real-world flexible docking scenarios.

As shown in S5, CoupleDiff achieves the lowest L-RMSD and the best reconstruction performance comparing to the other methods. According to b in S5, the drug engages in a complex network of interactions with key residues, including T380, M367, H422, V423, Q179, and L384, which collectively orchestrate the formation of a well-defined binding pocket, characterized not merely by static complementarity but by conformational plasticity, especially within the receptor's side-chain architectures. CoupleDiff demonstrates an understanding of this induced-fit paradigm by capturing the side-chain reorganization necessary for stable drug accommodation. In particular, CoupleDiff predicts a significant conformational shift in V423, which subtly reorients to alleviate steric hindrance and permit favorable van der Waals and hydrophobic contacts with Aceclofenac's chlorophenyl and phenylacetic moieties.

In contrast, FlexPose, despite achieving a comparable RMSD, fails to resolve a critical steric clash between V423 and the drug, which highlights the limitations of RMSD as a sole metric, as FlexPose's predicted pose superficially resembles the native

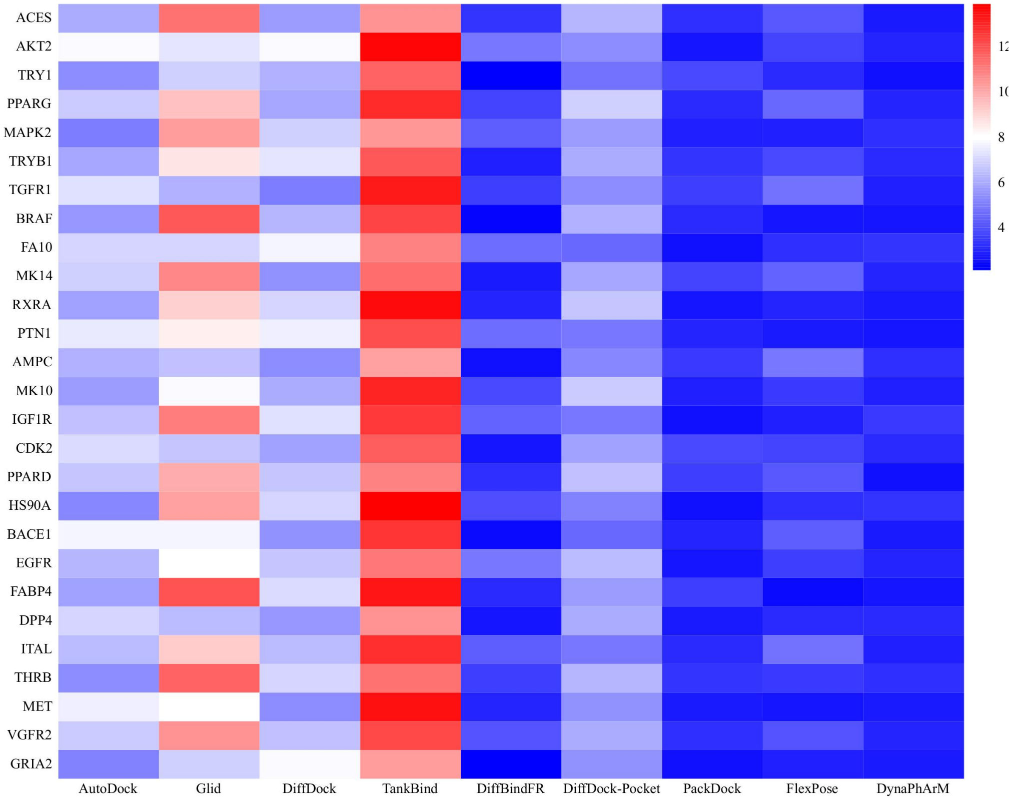


**Fig. S4.** Performance of different methods in reconstructing complex structures within the DUDE27 test set, evaluated by L-RMSD.

binding mode but lacks the microscopic interaction fidelity required for biological plausibility. The clash at V423 indicates FlexPose's insufficient modeling of side-chain entropy compensation.

Traditional metrics such as RMSD provide a coarse assessment of geometric alignment, they often obscure deeper deficiencies in interaction fidelity and physical plausibility. The above analysis reveals that accurately capturing side-chain flexibility and resolving steric conflicts—particularly at key residues like V423, which is essential for modeling biologically valid binding poses. CoupleDiff's superior performance stems not merely from pose prediction accuracy, but from its ability to model the energetic and structural consequences of induced fit at atomic resolution, which enables it to discriminate between superficially correct and mechanistically correct solutions—a capability critical for drug design applications where small structural inaccuracies can lead to large functional errors.

**J. Sensitivity Analysis of Priors.** To evaluate the model's robustness to potential inaccuracies in priors, we conducted a sensitivity analysis on the pre-computed Lennard-Jones (LJ) matrix. To measure the degradation in model performance, we added Gaussian noise with an increasing standard deviation (Noise Level) to the LJ matrix, with results summarized in Table S21.

**Table S21. Sensitivity analysis of the model to noise in the LJ prior. Performance is evaluated across different noise levels added to the LJ matrix.**

| Noise Level | Overall RMSD (Å) ↓ | sc-RMSD (Å) ↓ | Ligand Success Rate (<2Å) ↑ |
|---|---|---|---|
| 0.0 | 2.01 | 0.29 | 61.60% |
| 1.0 | 2.07 | 0.38 | 58.10% |
| 2.0 | 2.10 | 0.49 | 54.20% |
| 5.0 | 2.21 | 0.55 | 53.17% |
| No Prior | 2.11 | 0.59 | 52.91% |

At low noise levels, performance changed very little, as the model's learned data-driven patterns and the adaptive $\gamma_i$ compensated for minor inaccuracies. At very high noise levels, the prior became actively misleading. The performance degraded, but it did not collapse entirely. The performance plateaued at a level comparable to that of the model trained without any prior, demonstrating that the model could learn to effectively ignore a useless prior.

In conclusion, the model is not overly sensitive to errors in the AutoDock-generated LJ potential and can effectively mitigate its impact.

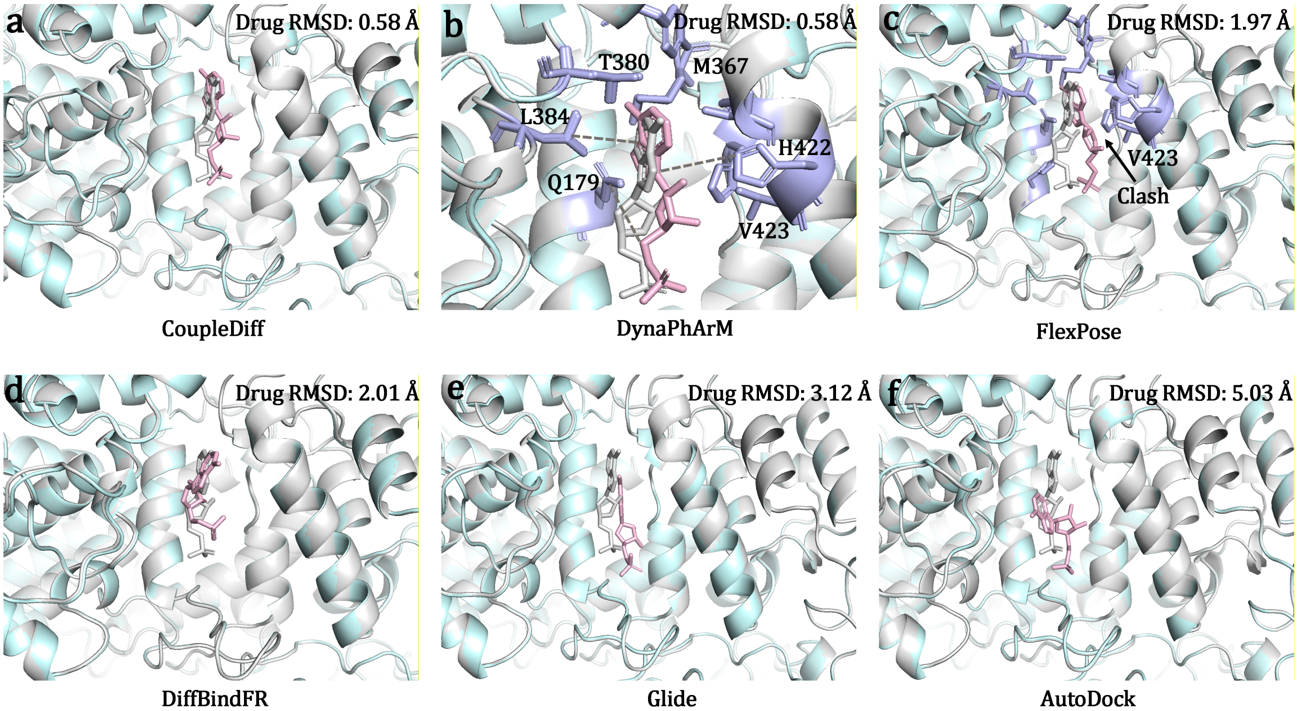


**Fig. S5.** The case of different methods on the reconstructing the structure of the complex which is consist of the drug Aceclofenac and the target 6Y3C from PDB. **a, d-f** The comparative figures illustrating CoupleDiff, DiffBindFR, Glide and AutoDock predicted structure (cyon target cartoon, pink drug sticks) alongside the experimental structure in grey from PDB, respectively. **b** The detailed illustration of a, which shows the key residues (purple) in the binding pocket and the interaction forces between key residues and drugs in dashed lines. **c** The comparative figures illustrating FlexPose (the second best method) predicted structure (cyon target cartoon, pink drug sticks) alongside the experimental structure in grey from PDB, which also identified key residues in purple.

### K. Structural Rationality Verification.

***K.1. Evaluation Metrics.*** The PoseBusters (PB) success rate reflects the proportion of predicted complexes that meet key biophysical and geometric criteria evaluated by the PB toolkit, including steric clashes, bond geometry, hydrogen bonding, and conformational strain energy, thereby excluding physically implausible poses.

***K.2. Experiments Results.*** Figure S6 clearly shows that while many methods can achieve a moderate success rate based on RMSD alone, they often fail the more stringent physical plausibility checks. CoupleDiff excels not only in geometric accuracy (RMSD $\leq 2$Å) but also demonstrates exceptionally high success rates (often $> 95\%$) across nearly all individual PB criteria, which indicates that the poses it generates are not only in the right place but are also chemically correct and free of high-energy conformations.

The main reasons are in its diffusion-based generative core tightly coupled with differentiable geometry and physics constraints. The traditional rigid docking only jumps between a few discrete conformations, making it easy to fall into the trap of just getting stuck in the pocket or atomic collision. The diffusion process is a continuous and smooth random walk in a high-dimensional coordinate space, with each small step iteration allowing the system to explore feasible surrounding areas and correct minor collisions or geometric deviations layer by layer. During each reverse-diffusion denoising step, the model explores a continuous manifold of drug poses, guided not only by statistical priors learned from known complexes but also by softly enforced vdW, hydrogen-bond, and geometric consistency losses. CoupleDiff naturally avoids steric clashes and spurious contacts, steering the generative trajectory toward low-energy, physically plausible basins. As a result, it attains high reconstruction fidelity, recapitulating native drug coordinates. This synergy between generative flexibility and physics-constrained regularization ensures CoupleDiff's poses consistently pass PB rigorous physical and geometric sanity checks.

### 4. Experimental Setting

The proposed CoupleDiff model is configured with a comprehensive set of hyperparameters covering feature embeddings, interaction modules, diffusion and decoder networks, physical constraints, and inference procedures. Table S23 summarizes the key settings, including embedding dimensions for target backbone, side-chain, and drug features, hidden sizes for multi-layer perceptrons in the interaction module, number of attention heads, and the output dimension of the physics-based interaction module feeding into the diffusion process. The diffusion module itself is trained over 1,000 timesteps with a denoiser network of hidden size 512, while the SE(3)-equivariant graph transformer decoder employs six layers with radial basis function (RBF) based message passing for coordinate refinement.

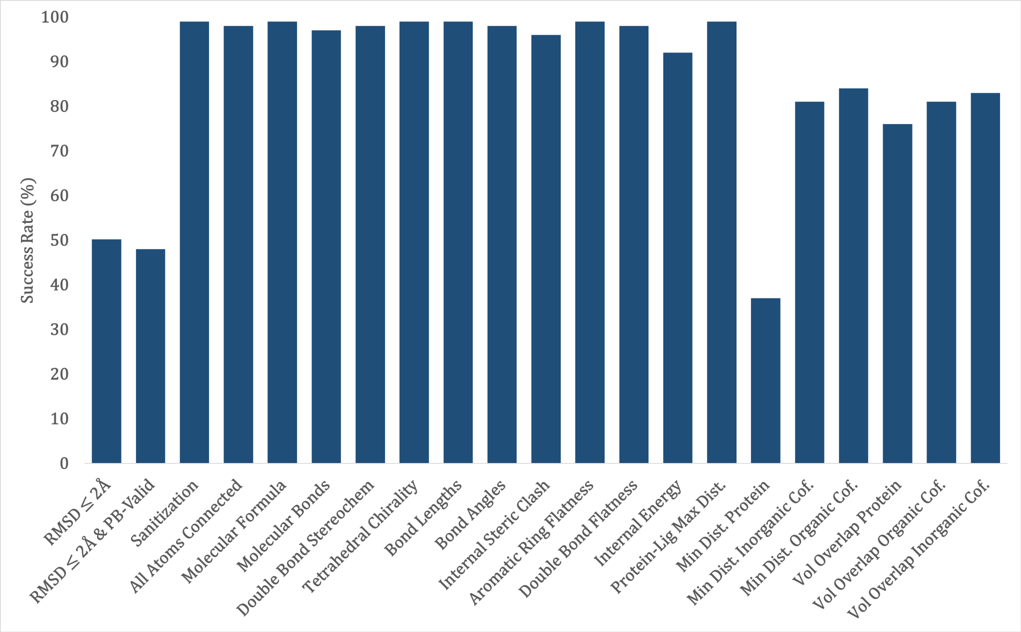


**Fig. S6.** Success rates of CoupleDiff generated poses across the 19 individual checks of the PB benchmark. Each bar represents the percentage of generated poses that successfully pass a specific physical or geometric validity check.

The optimization of the model's parameters is performed using the Adam algorithm, as implemented in the PyTorch library (27). The optimizer is applied to the learnable parameters of distinct architectural components, including the side-chain feature projector, the drug feature projector, the physics-based interaction module, the denoiser network within the diffusion module, and the SE(3)-equivariant graph transformer decoder. Adam computes individual adaptive learning rates for each parameter by utilizing estimates of the first and second moments of the gradients, which is particularly advantageous for a multifaceted model architecture such as the one employed, which integrates diverse modules requiring potentially different learning dynamics. The training process is initiated with a learning rate of $1 \times 10^{-4}$, while the $\beta_1$, $\beta_2$, and $\epsilon$ hyperparameters of the Adam optimizer are maintained at their default PyTorch values of 0.9, 0.999, and $1 \times 10^{-8}$, respectively.

**Table S23. Key hyperparameters used in the proposed model, covering embedding dimensions, interaction modules, diffusion and decoder configuration, physical constraints, and inference setup.**

| Name | Value | Description |
|---|---|---|
| D_E | 128 | Embedding dimension for backbone, side-chain, and drug features |
| D_K_INTERACTION | 128 | Key/Query/Value dimension |
| D_PRIME_INTERACTION | 256 | Output dimension of the physics-based interaction module, input to diffusion |
| MLP_HIDDEN_INTERACTION | 512 | Hidden dimension for MLP in the interaction module |
| N_ATTN_HEADS_INTERACTION | 4 | Number of attention heads |
| NUM_TRAIN_TIMESTEPS_DIFFUSION | 1000 | Total timesteps for the diffusion process during training |
| DENOISER_HIDDEN_DIFFUSION | 512 | Hidden dimension for the denoiser model |
| NUM_DECODER_LAYERS | 6 | Number of decoder layers |
| D_HIDDEN_ATTN_DECODER | 128 | Hidden dimension for attention in decoder |
| D_RBF_DECODER | 32 | Number of RBF kernels in decoder |
| D_MESSAGE_DECODER | 128 | Message dimension in decoder |
| D_MLP_HIDDEN_PHI_M_DECODER | 256 | Hidden dim. for MLP in message generation |
| D_MLP_HIDDEN_PSI_DECODER | 64 | Hidden dim. for MLP in coordinate update |
| LEARNING_RATE | 1e-4 | Learning rate |
| BATCH_SIZE | 64 | Batch size |
| NUM_EPOCHS | 50 | Number of epochs |
| REFINE_START_T_INFERENCE | 200 | Start timestep for denoising refinement |
| NUM_DENOISING_STEPS_INFERENCE | 50 | Steps in inference denoising loop |
| min_dist | 0.0 | Minimum distance for RBF |
| max_dist | 20.0 | Maximum distance for RBF |

Continued on next page

**Table S23 continued from previous page**

| Name | Value | Description |
|---|---|---|
| d_a_dist_max | 3.2 | Max donor–acceptor distance (H-bond) |
| d_a_dist_min | 2.5 | Min donor–acceptor distance (H-bond) |
| SIZE_X / SIZE_Y / SIZE_Z | 25 | Grid size in each spatial direction |
| NUM_MODES | 3 | Number of binding modes to generate |
| beta_start | 1e-4 | Start of noise schedule |
| beta_end | 0.02 | End of noise schedule |
| dropout_rate | 0.1 | Dropout rate |
| RMSD_THRESHOLD | 2.0 Å | Threshold for success (RMSD) |
| angle_min_deg | 120 | Minimum bond angle (H-bond) |
| angle_max_deg | 180 | Maximum bond angle (H-bond) |
| multiprocessing.num_workers | 64 | Number of worker processes for parallel tasks |
| multiprocessing.chunk_size | 10 | Chunk size for multiprocessing pool operations |
| drug_preparation.charge_model | gasteiger | Charge model used for ligand preparation |
| dataset_split.cd_hit_identity | 0.9 | Sequence identity threshold for CD-HIT clustering |
| dataset_split.validation_set_ratio | 0.2 | Proportion of data for the validation set |
| dataset_split.random_seed | 42 | Random seed for data splitting |
| backbone_encoder_params.num_layers | 4 | Number of layers in the backbone encoder |
| backbone_encoder_params.l_max_sh | 2 | Max degree for spherical harmonics |

Table S24 provides a comprehensive and detailed summary of all key settings, parameters, and protocols employed in the molecular dynamics (MD) simulations. It begins by specifying the software packages used for both protein and ligand modeling, along with the corresponding force fields, ensuring that interatomic interactions are represented with high fidelity and that the simulation parameters are fully compatible with the molecular systems under study. The table also details the system preparation procedures, including topology generation, file format conversion, solvation using explicit water models, and the addition of appropriate counterions to neutralize the system charge, thereby establishing a physically realistic and energetically balanced starting configuration suitable for subsequent dynamic simulations.

**Table S24. Molecular Dynamics Simulation Parameters**

| Parameter | Value |
|---|---|
| MD engine | GROMACS |
| Target parameterization | AmberTools (antechamber, parmchk2, tleap) |
| Topology conversion | ACPYPE |
| Target force field | amber99sb-ildn |
| Drug force field | GAFF |
| Drug charge model | Gasteiger (gas phase) |
| Water model | TIP3P |
| Box type | Cubic |
| Box distance from solute | 1.2 nm |
| Solvent model | spc216.gro |
| Ion addition | Neutralized with $Na^+$ and $Cl^-$ |
| Energy minimization algorithm | Steepest descent |
| Max minimization steps | 50,000 |
| EM convergence criterion | Fmax < 1000 kJ/mol/nm |
| Electrostatics method | PME |
| Coulomb cutoff | 1.0 nm |
| Van der Waals cutoff | 1.0 nm |
| Constraints | LINCS (all bonds) |
| NVT ensemble temperature | 300 K |
| NVT | V-rescale |
| NVT duration | 100 ps |
| Time step | 2 fs |
| NPT ensemble temperature | 300 K |
| Pressure | 1 bar |
| NPT | V-rescale |
| Barostat (equilibration) | Berendsen |
| Barostat (production) | Parrinello–Rahman |
| Compressibility | 4.5e-5 $bar^{-1}$ |
| NPT duration | 100 ps |
| Production simulation time | 500 ns |
| Trajectory output frequency | Every 10 ps |
| Energy output frequency | Every 10 ps |
| Periodic boundary conditions | 3D |
| Temperature coupling groups | Target__Drug and Water__and__ions |
| PBC correction | nojump and center |
| Target RMSD calculation | Backbone atoms |
| Drug RMSD calculation | Heavy atoms |
| Residue RMSF | Per-residue fluctuation |

The simulation workflow is described in a structured manner, covering energy minimization to remove unfavorable contacts, equilibration under both isothermal-isochoric (NVT) (constant number, volume, and temperature) and isothermal-isobaric (NPT) (constant number, pressure, and temperature) ensembles to stabilize the system thermodynamically, and production MD runs for collecting dynamic trajectories. Key parameters for each stage are listed, including integration time steps, total simulation durations, thermostat and barostat algorithms, target temperature and pressure values, and the application of periodic boundary conditions to mimic bulk solvent behavior.

Furthermore, the table specifies parameters related to trajectory and energy output, such as sampling frequencies, which are essential for accurate monitoring of structural evolution, and any applied positional restraints or bond constraints used to maintain critical structural features. It also highlights post-processing and analysis procedures, including calculation of RMSD and RMSF values, which allow quantitative assessment of the stability, flexibility, and conformational dynamics of the target-drug complexes. Collectively, Table S24 provides an exhaustive overview of the computational setup, encompassing preparation, execution, and analysis, to ensure rigorous reproducibility, accuracy, and reliability of the MD simulation results.

### 5. Extended Materials and Methods

**Model Validation and Evaluation.** The proposed method evaluates both structural and functional performance. For structural accuracy, we compute the root mean square deviation (RMSD) of the entire protein–ligand complex (overall RMSD), the ligand's 3D coordinates (L-RMSD), and the side-chain conformations (sc-RMSD). Corresponding success rates are reported as the percentage of cases with RMSD values less than 2 Å, including overall success rate, ligand success rate, and side-chain success rate, which respectively reflect the reconstruction fidelity of the full complex, the ligand molecule, and side-chain positioning. To further evaluate conformational flexibility and dynamic stability, we compute the root mean square fluctuation (RMSF) per residue.

**Implementation and Training Details.** Our model, CoupleDiff, and the entire data processing pipeline were implemented in Python using the PyTorch deep learning framework. All computationally intensive tasks, including model training and LJ matrix generation, were conducted on a single NVIDIA GeForce RTX 4090 GPU, with data parallelism managed by 64 worker processes.

Our workflow begins with a multi-stage data preprocessing pipeline. Protein and ligand structures from sources like PDBbind and CASF-2016 were first prepared into the PDBQT format using MGLTools (28) with the Gasteiger charge model. To provide a standardized starting pose for our refinement model, we performed an initial coarse docking using AutoDock Vina (5), configured with an exhaustiveness of 16. Following this, each complex was split into its constituent backbone, side-chain, and ligand components, from which we generated a comprehensive set of features. Graph representations were constructed for each component, encoding detailed chemical and geometric properties as described in Supplementary Section A.1. Crucially, we pre-computed an LJ interaction matrix for every complex to inform our physics-constrained attention module, alongside ground-truth labels for our physics and geometry-guided loss.

The architecture of CoupleDiff is defined by several key components. The encoders for the protein backbone, side chains, and the ligand each consist of four layers. The interaction module utilizes four attention heads with an embedding dimension of 128. The diffusion process is configured with 50 timesteps following a linear noise schedule, and the denoiser network uses a hidden dimension of 512. The final SE(3)-equivariant decoder is composed of six layers. A complete list of all hyperparameters is provided in Supplementary Section D.

The model was trained end-to-end using the Adam optimizer with a learning rate of $1 \times 10^{-4}$ and a weight decay of $1 \times 10^{-5}$. We employed a batch size of four and trained for a maximum of 200 epochs, with an early stopping criterion based on validation performance. A key innovation in our training strategy is the use of dynamic weighting for the multi-task loss function. This strategy adaptively balances the contributions from the structure reconstruction, diffusion denoising, and physics-guided loss components, ensuring a stable and balanced convergence across all objectives as detailed in Supplementary Section C.7. For all comparative experiments, we followed established data splitting protocols, using a 90% sequence identity threshold for CD-HIT to create non-redundant training and validation sets, and all reported metrics are the average of multiple runs to ensure statistical reliability.

**Quantification of Steric Clashes.** Steric clashes were quantified based on the spatial overlap between atomic van der Waals spheres. For each pair of atoms, the Euclidean distance between their atomic centers was compared with the sum of their theoretical vdW radii. To account for physically allowable atomic overlap and uncertainties associated with structural coordinates, a tolerance factor $\tau$ was applied to the pairwise vdW distance threshold.

For a ligand containing $N_L$ atoms and a protein containing $N_P$ atoms, let $d(i, j)$ denote the Euclidean distance between atoms $i$ and $j$, and let $R_i$ and $R_j$ denote their corresponding vdW radii. The number of intramolecular steric clashes within the ligand was calculated as

$$C_{\text{intra}} = \sum_{i=1}^{N_L} \sum_{j=i+1}^{N_L} \mathbb{I}\left[d(i,j) < \tau(R_i + R_j)\right] \cdot \mathbb{I}\left[(i,j) \notin E_{\text{cov,det}}\right],$$

where $E_{\text{cov,det}}$ denotes the set of ligand atom pairs that are covalently connected or separated by fewer than three bonds. These atom pairs were excluded because their spatial proximity is determined by molecular connectivity rather than unfavorable steric overlap.

Intermolecular steric clashes between ligand and protein atoms were calculated as

$$C_{\text{inter}} = \sum_{i=1}^{N_L} \sum_{k=1}^{N_P} \mathbb{I}\left[d(i,k) < \tau(R_i + R_k)\right].$$

Here, $\mathbb{I}[\cdot]$ denotes the indicator function, which equals one when the specified condition is satisfied and zero otherwise. Thus, $C_{\text{intra}}$ and $C_{\text{inter}}$ represent the total numbers of intramolecular and intermolecular atom pairs, respectively, whose separation falls below the tolerated vdW distance threshold.

**All-atom Molecular Dynamics Simulation Protocol.** MD simulations were performed using GROMACS (29). For the CAII analysis, the holo complex used as the starting structure for MD was the conformation generated by CoupleDiff from the apo receptor input, and the apo receptor was simulated separately as a reference; production simulations were run for 500 ns per system. The workflow included protein–ligand structure preparation, topology construction, system solvation and ionization, sequential equilibration, and long-timescale production simulation. Detailed simulation parameters are summarized in Supplementary Section D.

The docked protein–ligand complex was first separated into protein and ligand components for independent topology generation. The ligand was parameterized using the general Amber force field (30). Atomic charges were assigned using the Gasteiger method via Antechamber, and missing bonded and nonbonded parameters were generated using parmchk2. AMBER-compatible topology and coordinate files were constructed using tleap and subsequently converted into GROMACS-compatible formats using acpype. The protein was described using the amber99sb-ildn force field (31), and its topology was generated using the pdb2gmx module. Explicit solvent molecules were modeled using the TIP3P water model (32). During equilibration, position restraints were applied to both the protein and the ligand to preserve the initial docked configuration while allowing solvent relaxation.

The protein–ligand complex was embedded in a periodic cubic simulation box with buffer space to prevent self-interaction across periodic boundaries. The system was solvated with explicit water molecules and neutralized by adding appropriate counterions. All topology components, including force field definitions and drug parameters, were integrated to ensure consistent treatment of bonded and nonbonded interactions throughout the system.

Before production simulation, the system underwent a multi-stage relaxation protocol. Energy minimization was performed to eliminate unfavorable steric contacts and relieve local strain. This was followed by canonical ensemble equilibration to stabilize the system temperature, and isothermal-isobaric ensemble (NPT) equilibration to achieve proper system density and pressure stabilization. Temperature and pressure were controlled using established thermostat and barostat schemes to ensure stable ensemble conditions.

Subsequently, long-timescale production MD simulations were conducted under NPT conditions. Periodic boundary conditions were applied in all spatial dimensions, long-range electrostatic interactions were treated using the particle mesh Ewald method (33), and covalent bonds involving hydrogen atoms were constrained to enable stable integration with a femtosecond time step. The resulting trajectories were used to evaluate the dynamic stability of the protein–ligand complexes and to characterize interaction patterns over time.

**Data Collection and Preprocessing.** We curated a dataset of 21,762 high-quality protein–ligand complexes from the Protein Data Bank (PDB). To prevent data leakage and ensure generalizability, complexes were grouped by 30% sequence identity, with entire clusters assigned to either the training (60%), validation (20%), and test sets (20%), guaranteeing no sequence overlap between subsets. The details of the training set can be found in Supplementary Section B.

For the complex structure reconstruction task, we assessed the performance of CoupleDiff to regenerate known binding poses using four distinct benchmarks. These include the CASF2016-Drug dataset, a refined subset of the widely used CASF2016 benchmark (34), focusing exclusively on drug-like ligands and their corresponding protein targets, emphasizing biologically and pharmacologically meaningful interactions, and providing a more targeted benchmark for evaluating models in drug–target docking; the Astex set, a widely-used high-quality benchmark comprising 85 protein–ligand complexes known for its chemical diversity and drug-like ligands, providing a robust measure of performance on pharmacologically relevant complexes (35); and the ESMFold-predicted set built upon receptor structures generated by the ESMFold (36) protein structure prediction model, to evaluate performance in more realistic scenarios where experimental structures are unavailable, thereby testing the robustness of the proposed model to structural inaccuracies. Finally, beyond positional accuracy, we utilized the PB dataset (37). The PB dataset and toolkit form a comprehensive benchmark suite specifically designed to evaluate the structural plausibility and physical validity of protein–ligand complex models, which includes 428 high-quality crystal structures of protein–ligand complexes. By highlighting not only how close a predicted pose is, but also whether it makes chemical and physical sense. The detailed results on PB dataset can be found in Supplementary Section C.11.

In our evaluation of CoupleDiff's performance on the cross docking task, we compared it with a variety of methods using several benchmark datasets, including CDK2, EGFR, DUDE27-HoloEns, and ANN. The CDK2 and EGFR datasets are curated collections of high-resolution, inhibitor-bound protein–ligand complexes from the PDB (38). The CDK2 set includes 11,317 Cyclin-Dependent Kinase 2 complexes, emphasizing ATP-competitive interactions in both rigid and flexible kinase active sites. EGFR comprises 67 wild-type and mutant kinase-inhibitor pairs, challenging algorithms to accommodate subtle shifts in the activation loop and $\alpha$C-helix. The DUDE27-HoloEns dataset (39) is a curated ensemble-docking benchmark comprising 27 DUD-E targets for which multiple holo structures have been experimentally determined. Each target's ensemble includes all available high-resolution co-crystal complexes, yielding 268 pairs of complexes. The ANN dataset (40) comprises 65 protein–ligand systems, each represented by a native holo complex and one or more non-native receptor conformations into which the cognate ligand is re-docked. Designed specifically for cross-docking, it challenges algorithms to recover the correct binding pose despite backbone and side-chain rearrangements not present in the ligand's original crystal structure, thereby gauging a method's robustness to receptor conformational variability.
